# iTAP, a novel iRhom interactor, controls TNF secretion by policing the stability of iRhom/TACE

**DOI:** 10.1101/255745

**Authors:** Ioanna Oikonomidi, Emma Burbridge, Miguel Cavadas, Graeme Sullivan, Danielle Clancy, Blanka Collis, Jana Brezinova, Jana Humpolickova, Tianyi Hu, Andrea Bileck, Christopher Gerner, Alfonso Bolado, Alex von Kriegsheim, Kvido Strisovsky, Seamus J. Martin, Colin Adrain

## Abstract

The apical inflammatory cytokine TNF regulates numerous important biological processes including inflammation and cell death, and drives inflammatory diseases. TNF secretion requires ADAM17/TACE, which cleaves TNF from its transmembrane tether, releasing it for signalling. The trafficking of ADAM17/TACE to the cell surface, and stimulation of its proteolytic activity, depends on membrane proteins, called iRhoms. To delineate how the TNF/TACE/iRhom axis is regulated, we performed an immunoprecipitation/mass spectrometry screen to identify iRhom-binding proteins. Here we report a novel protein, that we name iTAP (iRhom tail-<u>a</u>ssociated <u>p</u>rotein) that binds to iRhoms, enhancing the stability of iRhoms and TACE, preventing their degradation in lysosomes. iTAP-null primary human macrophages, or tissues from iTAP KO mice, are dramatically depleted in the levels of iRhom2 and active TACE, and are, consequently, profoundly impaired in TNF production. Our work illustrates iTAP as a physiological rheostat controlling TNF signalling and a novel target for the control of inflammation.

## Introduction

The cytokine TNF controls numerous important biological processes (e.g. inflammation, fever, apoptosis, necroptosis, cachexia, tumorigenesis, viral infection, insulin signaling) and is heavily implicated in inflammatory disease (Brenner et al., 2015). Anti-TNF biologics are the highest-selling drugs internationally and there is intense interest in how TNF secretion is regulated (Brenner et al., 2015).

TNF is synthesized with a transmembrane tether, but much of its biology is driven by its secreted form, whose release from the cell surface is catalyzed by the protease TACE (<u>T</u>NF <u>α</u> <u>c</u>onverting <u>e</u>nzyme) (Horiuchi et al., 2007; Peschon et al., 1998), also called ADAM17 (<u>a</u> <u>d</u>isintegrin <u>a</u>nd <u>m</u>etalloprotease) (Gooz, 2010; Zunke and Rose-John, 2017). Other prominent TACE substrates include the activating ligands of the EGFR (<u>e</u>pidermal <u>g</u>rowth factor receptor), an important pathway that drives growth control.

Given its ability to elicit potent biological responses, it is unsurprising that TACE is stringently regulated (Murphy, 2009; Grötzinger et al., 2017). A major control point in TACE regulation involves its trafficking within the secretory pathway (Schlondorff et al., 2000). TACE is synthesized in the endoplasmic reticulum (ER) as a catalytically inactive precursor. For TACE to become proteolytically active, it must undergo a maturation step—removal of its prodomain— which is catalyzed by proprotein convertases in the *trans*-Golgi (Schlondorff et al., 2000). The exit of TACE from the ER and its trafficking to the cell surface requires regulatory proteins called iRhoms (Adrain et al., 2012; McIlwain et al., 2012; Li et al., 2015). Hence, iRhom KO mice or cells lack TACE activity (Adrain et al., 2012; Christova et al., 2013; Li et al., 2015). Mice null for iRhom2, whose expression is enriched in myeloid cells, cannot secrete TNF (Adrain et al., 2012; McIlwain et al., 2012; Siggs et al., 2012).

An important checkpoint to license TACE activity involves its stimulation by agents including phorbol esters, Toll-like receptor agonists and G-protein coupled receptor ligands (Grötzinger et al., 2017; Arribas et al., 1996; Hall and Blobel, 2012; Brandl et al., 2010; Wetzker and Böhmer, 2003). iRhoms play a central role in this cell surface stimulation mechanism. TACE-activating stimuli provoke the phosphorylation of the cytoplasmic tail of iRhom2, leading to the recruitment of 14-3-3 proteins, enforcing the detachment of TACE from iRhom2, and facilitating its ability to cleave TNF (Grieve et al., 2017; Cavadas et al., 2017). Hence, the trafficking fate of iRhoms and TACE is coupled until TACE stimulation on the cell surface, when the proteins dissociate and presumably assume different fates.

In spite of the importance of trafficking for TACE regulation (Schlondorff et al., 2000; Adrain et al., 2012; Dombernowsky et al., 2015), surprisingly little is known about the machinery that controls TACE trafficking to/from the plasma membrane. An exception is PACS-2 (<u>p</u>hosphofurin <u>a</u>cidic <u>c</u>luster <u>s</u>orting protein <u>2</u>), a protein that colocalizes with mature TACE in endocytic compartments (Dombernowsky et al., 2015) and controls its endocytic recycling. Ablation of PACS-2 in cells impairs the cell surface availability of TACE, reducing substrate cleavage (Dombernowsky et al., 2015). However, PACS-2 has a relatively modest impact on TACE biology *in vivo* (Dombernowsky et al., 2017), suggesting the possibility of unidentified trafficking regulators that may act separately from, or redundantly with, PACS-2.

As iRhoms form functionally important complexes with cell surface TACE and are endocytosed and degraded in lysosomes, modulation of iRhom endocytic trafficking has the potential to act as a regulatory mechanism to control TNF secretion. However, the machinery involved in maintaining stable cell surface levels of iRhoms is unknown. Here we identify a novel protein that we name iTAP (iRhom tail-<u>a</u>ssociated <u>p</u>rotein) that is essential for the control of the stability of iRhom2 and TACE in the late secretory pathway. Ablation of iTAP triggers the mis-sorting of iRhom and mature TACE to lysosomes, where they are degraded. Consistent with this, loss of iTAP results in a dramatic reduction in TACE activity and TNF secretion. Our work reveals iTAP as a key physiological regulator of TNF release.

## Results

### iTAP, a novel interactor of iRhoms, is an atypical FERM domain-containing protein

To identify novel regulators of mammalian iRhoms 1 and −2, we adopted an immunoprecipitation/mass spectrometry (IP/MS) approach similar to the one that was validated in our previous work (Cavadas et al., 2017). As shown in Figure 1A, we generated a panel of HEK 293ET cell lines stably expressing HA-tagged forms of full-length iRhom1, iRhom2, or the iRhom1 N-terminal cytoplasmic tail only. To focus only on proteins that bind selectively to iRhoms, we included several related rhomboid-like proteins as specificity controls (Fig. 1A). As expected, only immunoprecipitates (IPs) from cells expressing full-length HA-tagged iRhom1 or iRhom2 captured endogenous TACE, confirming the validity of the approach (Fig. 1B).

**Figure 1.**
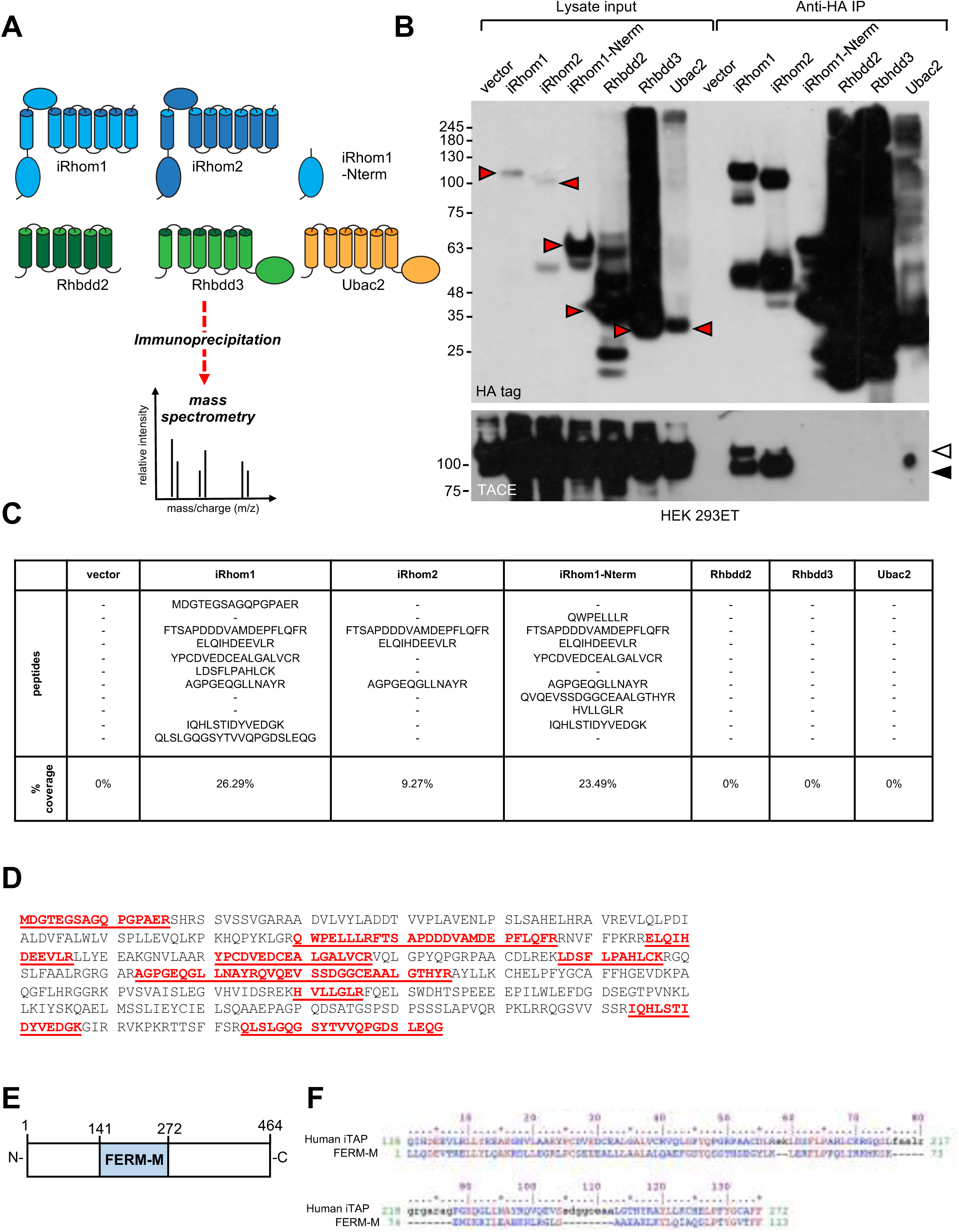
Identification of iTAP as a novel iRhom interacting protein. (**A**) Schematic diagram showing the stable HEK 293ET cell lines expressing iRhom proteins or related rhomboid pseudoproteases as controls, which were subjected to immunoprecipitation followed by mass spectrometry. (**B**) An example immunoprecipitation from the cell lines indicating that only immunoprecipitates from cell lines expressing WT iRhom1 or iRhom2 contain the binding of the positive control protein, TACE. Here and throughout, immature and mature species of TACE are indicated with white and black arrowheads respectively. The red arrows show the full-length forms of the individual rhomboid-like proteins. (**C**) Peptides assigned to FRMD8 that were found in immunoprecipitates from iRhom1, iRhom2 or the N-terminus of iRhom1 but not in the other samples. The peptides are shown from a representative experiment. (**D**) All of the peptides found in the iRhom immunoprecipitates from (C) were mapped unto the human FRMD8 amino acid sequence. (**E**) Schematic diagram illustrating the domain structure of iTAP/FRMD8. (**F**) Alignment of the central FERM lobe (FERM-M) from iTAP/FRMD8 versus the pfam FERM-M consensus (<u>pfam00373</u>).

To identify novel iRhom-binding proteins, we subjected IPs from these cells to mass spectrometry. This analysis revealed, in multiple replicate experiments, peptides from a largely uncharacterized protein, called FRMD8, in IPs of iRhom1 or iRhom2, but not in control IPs (Fig. 1C,D). Furthermore, FRMD8 was found in IPs from cells expressing the N-terminus of iRhom1 (Fig. 1A,C), suggesting that it was recruited to the iRhom cytoplasmic tail (R-domain) of iRhom, an important regulatory region (Cavadas et al., 2017; Grieve et al., 2017; Hosur et al., 2014; Maney et al., 2015). In light of this, we named the novel protein iTAP (‘iRhom tail <u>a</u>ssociated <u>p</u>rotein’). A closer inspection of the iTAP sequence revealed that it encodes a FERM (band <u>4</u>.1/<u>e</u>zrin/radixin/<u>m</u>oesin) domain (Chishti et al.) (Fig. 1E,F). Proteins containing FERM domains fulfil many important roles, including signaling, and organization of the cell cortex(Fehon et al., 2010; Hoover and Bryant, 2000; Baines et al., 2014; Moleirinho et al., 2013). The well-characterized FERM domain contains three distinct lobes that together resemble a three-leaved clover (Pearson et al., 2000). However, unlike most FERM domain-containing proteins, but similar to its paralog KRIT1—an adaptor protein in the cerebral cavernous malformation pathway that regulates the establishment of vasculature(Pal et al., 2017), iTAP contains only the central (FERM-M) lobe. iTAP orthologs are present in metazoans, including *Drosophila* and *Danio* (Kategaya et al., 2009). The iTAP protein is expressed broadly and is co-expressed with TACE and iRhom1 or iRhom2 in a range of tissues relevant for TACE biology (Fig. S1A,B).

Independent immunoprecipitation experiments verified that iTAP binds specifically to both iRhom1 and iRhom2, but not to the related rhomboid pseudoproteases Ubac2 and Rhbdd2 (Fig. 2A). iTAP binds the cytoplasmic tail of iRhoms, since a mutant containing only the cytoplasmic tail of iRhom1 bound iTAP robustly, whereas a mutant lacking all of the iRhom2 cytoplasmic tail (∆Nterm) failed to bind (Fig. 2A). By contrast, removal of the iRhom homology domain (IRHD), the luminal globular domain between transmembrane helices 1 and 2 (Fig. 2B) from iRhom2 had no impact on iTAP recruitment (Fig. 2A). These data indicate that iTAP is specifically recruited to the cytoplasmic tail of iRhoms. Notably, when iTAP-FLAG, but not a panel of control proteins, was immunoprecipitated from cell lysates, we detected the binding of endogenous iRhom1 and iRhom2 to iTAP (Table S1). These data confirm that iTAP is a specific endogenous interactor of iRhoms.

**Figure 2.**
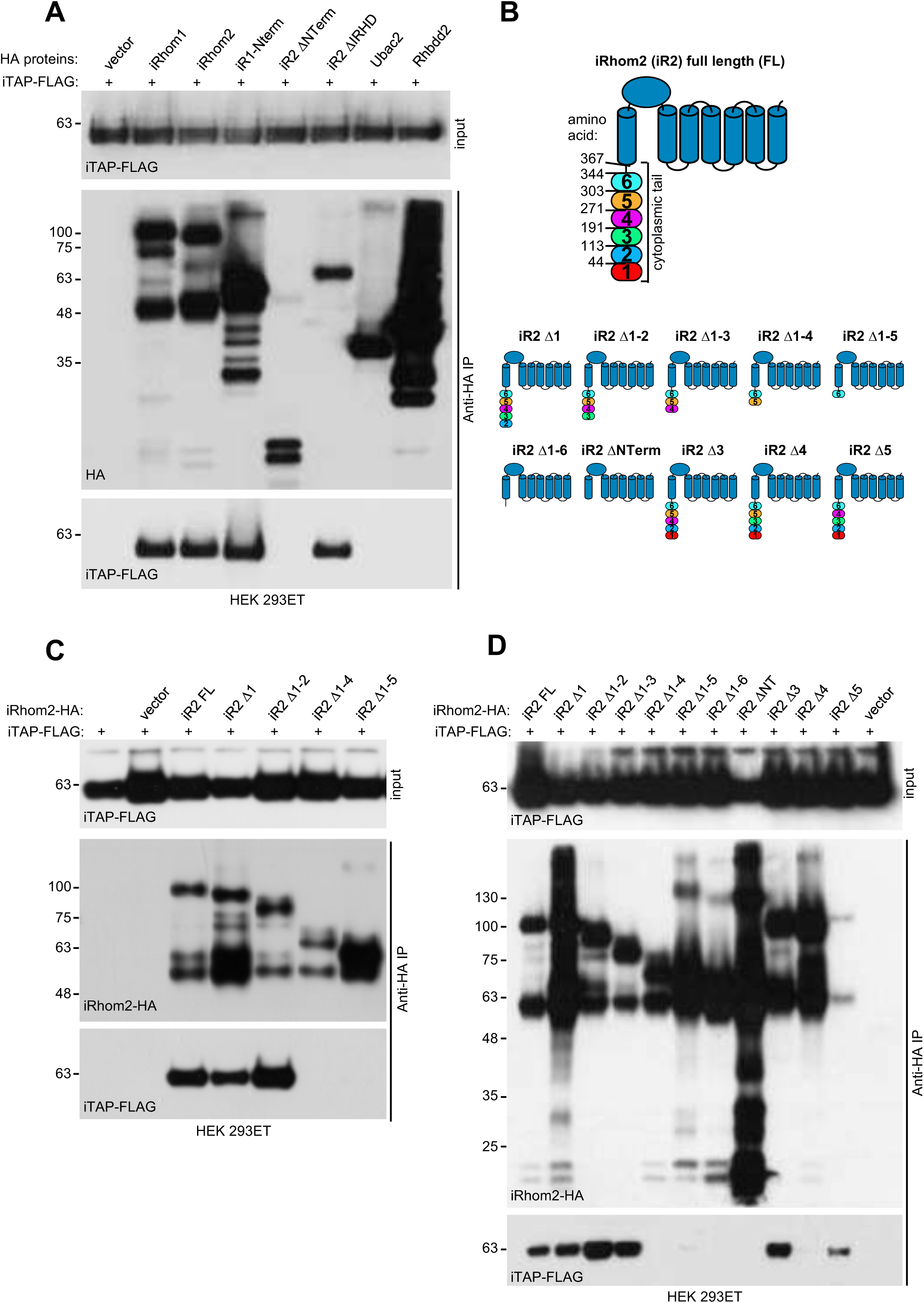
Validation of iTAP binding to iRhom2. (**A**) iTAP binds specifically to iRhom −1 and −2. Human iTAP-FLAG was transfected into HEK 293ET cells together with empty vector or the indicated HA-tagged iRhoms, their deletion mutants or the rhomboid pseudoproteases Ubac2 or Rhbdd2. HA proteins were immunoprecipitated and FLAG binding was assessed by western blot. (**B**) Schematic diagram indicating truncation mutants of the iRhom2 cytoplasmic tail that were generated to map the iTAP binding region. The cytoplasmic tail of iRhom2 was divided into 6 arbitrary portions. (**C,D**) iTAP binds to iRhom2 within an area defined by region 4 of iRhom2 (192-271aa). HEK 293ET cells were transfected with iTAP-FLAG and iRhom2-HA full length (FL) or the indicated iRhom2-HA deletion constructs shown in (B) Anti-HA immunoprecipitates were assessed for the binding of iTAP-FLAG by western blotting.

To delineate the region within the cytoplasmic tail of iRhom that iTAP binds to, we created a series of sequential truncations, or deletions, of the iRhom2 cytoplasmic tail (Fig. 2B). This revealed that amino acids 191-271 of the iRhom2 tail contain the main determinant for iTAP binding (Fig. 2C,D).

We next examined the cellular localization of GFP-tagged iTAP in fixed and permeabilized mammalian cells. As shown in Figure 3A, iTAP-GFP exhibited a powdery staining in the cytoplasm and nucleus. When co-expressed with mCherry-tagged iRhom2, iTAP was recruited to iRhom2, which is itself predominantly found in the ER under these overexpression, fixation and permeabilization conditions (Zettl et al., 2011) (Fig 3B).

**Figure 3.**
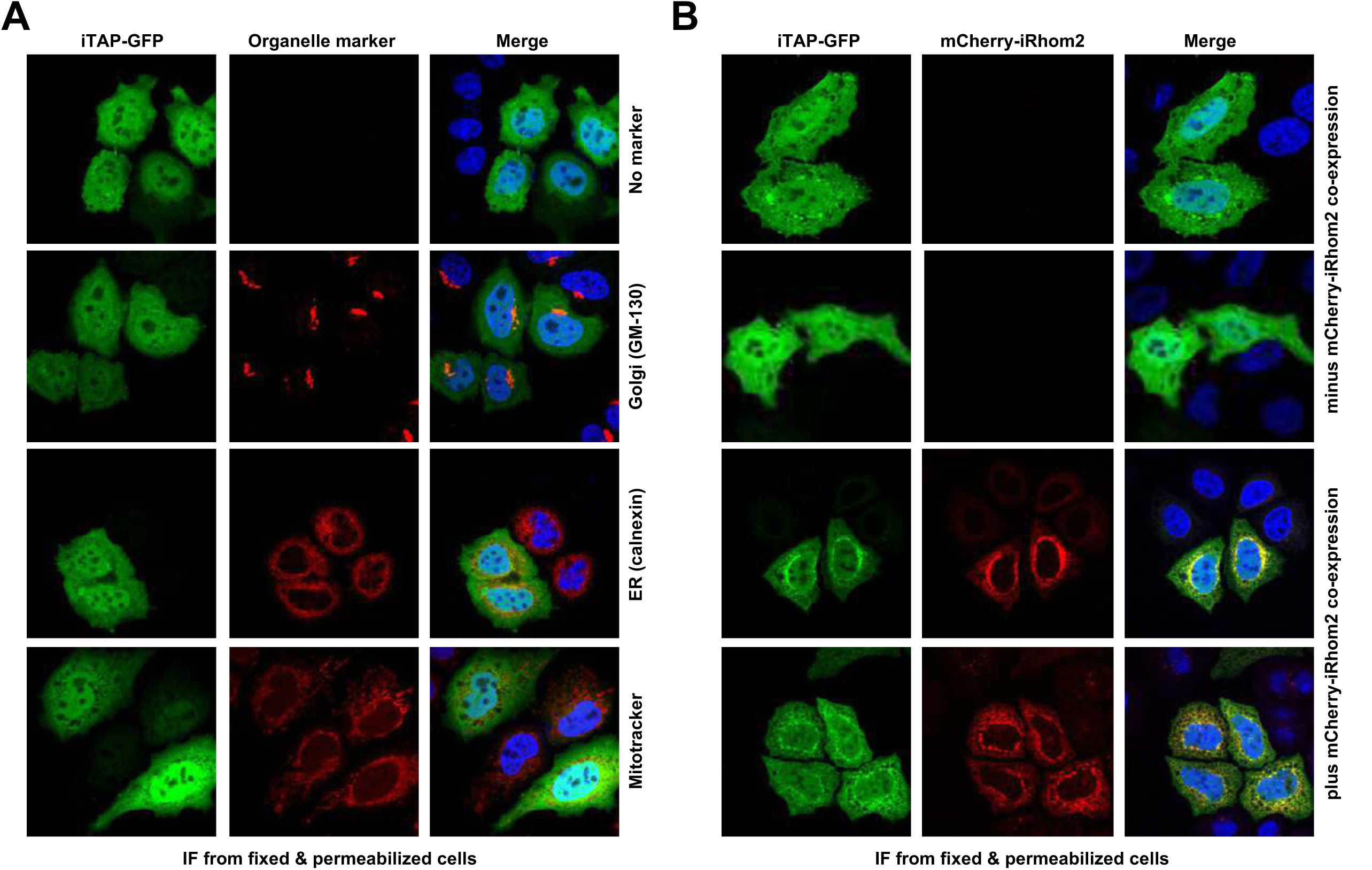
Cellular localization of iTAP. (**A**) HeLa cells were transfected with iTAP-GFP (600ng). After 24 h, cells were fixed and permeabilised and immunostained, as indicated, with specific antibodies against Golgi (GM-130) or ER (Calnexin). The fluorescent dye Mitotracker-Red was used to visualise mitochondria. DNA was stained with Hoescht (blue). (**B**) HeLa cells were transfected with iTAP-GFP (600 ng) alone, or in the presence of iRhom2-Cherry (600ng), as indicated. After 24 h, cells were fixed and stained with Hoescht to visualise DNA (blue). Cells were mounted and visualised by confocal microscopy.

### iTAP-deficient cells have impaired TACE shedding activity

To determine the functional importance of iTAP binding to iRhom, we used CRISPR to ablate iTAP in HEK 293ET cells, which was confirmed by the lack of iTAP protein expression (Fig. 4A). As TACE trafficking and cell surface stimulation depends on iRhoms, we examined the ability of WT versus iTAP-null cells to support TACE sheddase activity. Notably, the PMA-induced shedding of a panel of chimeric alkaline phosphatase (AP) TACE substrates (Sahin et al., 2004), including EGFR ligands and TNF, was substantially impaired in *iTAP* KO cells (Fig. 4B).

**Figure 4.**
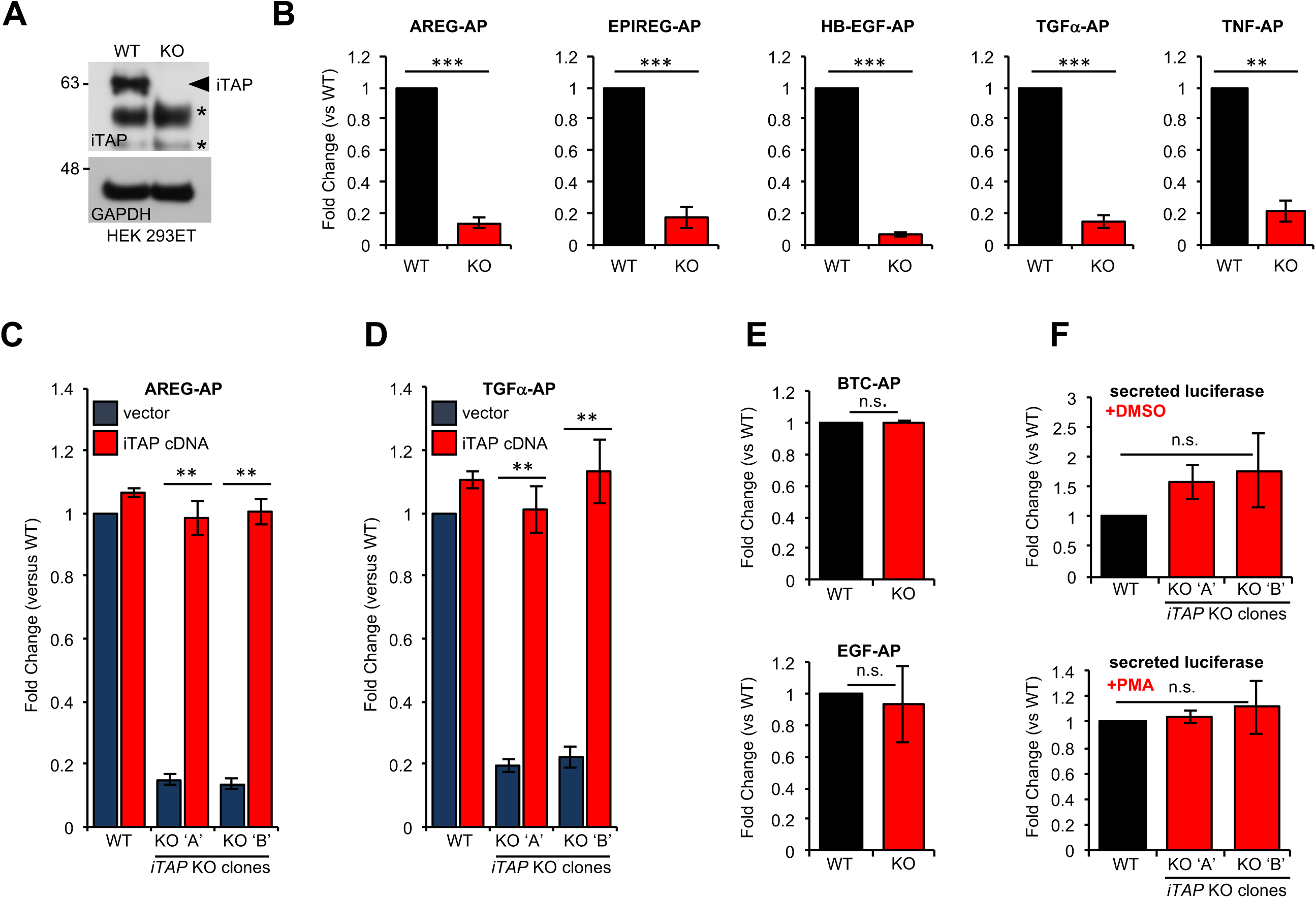
KO of iTAP diminishes TACE proteolytic activity. (**A**) Anti-iTAP immunoprecipitates from WT versus iTAP KO HEK 293ET cells were analyzed by immunoblotting. A GAPDH blot is the loading control for the inputs. **(B)** PMA-stimulated TACE shedding is impaired in iTAP KO cells. The TACE substrates [Amphiregulin (AREG), Epiregulin (EPIREG), Heparin Binding-Epidermal Growth Factor (HB-EGF), Transforming Growth Factor-α (TGFα) and Tumor Necrosis Factor (TNF)] fused to alkaline phosphatase (AP) were transfected into HEK 293ET WT or iTAP KO cells. TACE activity was assessed based on AP activity secreted into the supernatant of the cells as described in materials and methods. (**C,D**) Expression of iTAP rescues the impaired shedding in iTAP KO cells. WT or iTAP KO HEK 293ET cells stably expressing empty vector or human iTAP were transfected with AREG-AP or TGFα-AP, then challenged in PMA shedding assays as described above. (**E**) The shedding impairment is specific to TACE. ADAM10 AP-fused substrates [Betacellulin (BTC) and Epidermal Growth Factor (EGF)] were transfected into the WT vs iTAP KO HEK 293ET cells. The cells were treated with the ADAM10 stimulant Ionomycin (IO) and AP activity was measured in the medium. (**F**) Global secretion is not impaired in iTAP KO cells. WT or iTAP KO cells were transfected with secreted luciferase and luciferase associated luminescence was measured in the supernatant of vehicle (DMS0, upper graph) or PMA-stimulated cells (lower graph). Here and throughout: KO ‘A’ and KO ‘B’ denotes independent iTAP KO HEK 293ET clones. PMA (1 μM) or IO (2.5 μM) incubations took place for 1h following serum starvation. Shedding or secretion values are expressed as fold change relative to WT cells. Data are presented as mean ± standard deviation and represent at least 3 independent experiments. * = p≤0.05, ** = p≤0.01, *** = p≤0.001 and n.s. = non-significant.

This shedding defect was rescued by the expression of an *iTAP* cDNA, confirming that the loss of iTAP was directly responsible for defective TACE activity (Fig.4C,D). By contrast, the cleavage of the EGFR ligands EGF and BTC, which are cleaved specifically by ADAM10, the metalloprotease most closely related to TACE, was unaffected (Fig. 4E). The release of a model secreted substrate was unimpaired in iTAP KO cells (Fig. 4F) confirming that iTAP is a highly specific regulator of the TACE pathway, but does not affect secretion in general.

### Proteolytically-active TACE levels are specifically depleted in iTAP-deficient cells

To investigate how loss of iTAP affected TACE, we examined the maturation status of TACE, a readout for its trafficking and activation status (Adrain et al., 2012). Strikingly, iTAP-null cell lines (Fig. 5A) showed a dramatic depletion of mature TACE, which is identified by its faster migration pattern (Fig. 5B). To confirm this, we treated lysates with the deglycosylating enzymes Endoglycosidase-H (Endo-H), which removes high mannose N-linked glycans added in the ER, but not complex N-linked glycans found in the later secretory pathway, versus PNGase F, which deglycosylates both (Fig. 5C). This analysis confirmed that iTAP KO cell lines were dramatically depleted of mature TACE. Interestingly, this is a similar, but less penetrant, phenotype to that found in iRhom1/2 double knockout (DKO) MEFs (Christova et al., 2013), since residual levels of TACE remained in iTAP-null cells. Notably, mature TACE could be rescued by iTAP overexpression in iTAP KO cells, confirming the specificity of the effect (Fig. 5D). Overexpression of iTAP in WT cells also modestly enhanced mature TACE levels (Fig. 5D). Biotinylation experiments using a non-cell permeable cross-linker, revealed that iTAP KO cells had substantially depleted mature TACE levels on the cell surface (Fig. 5E), explaining the basis of the TACE activity defect. The pronounced impact of iTAP on TACE maturation was highly specific for TACE, since the maturation of other related ADAM proteases was unaffected (Fig. 5F).

**Figure 5.**
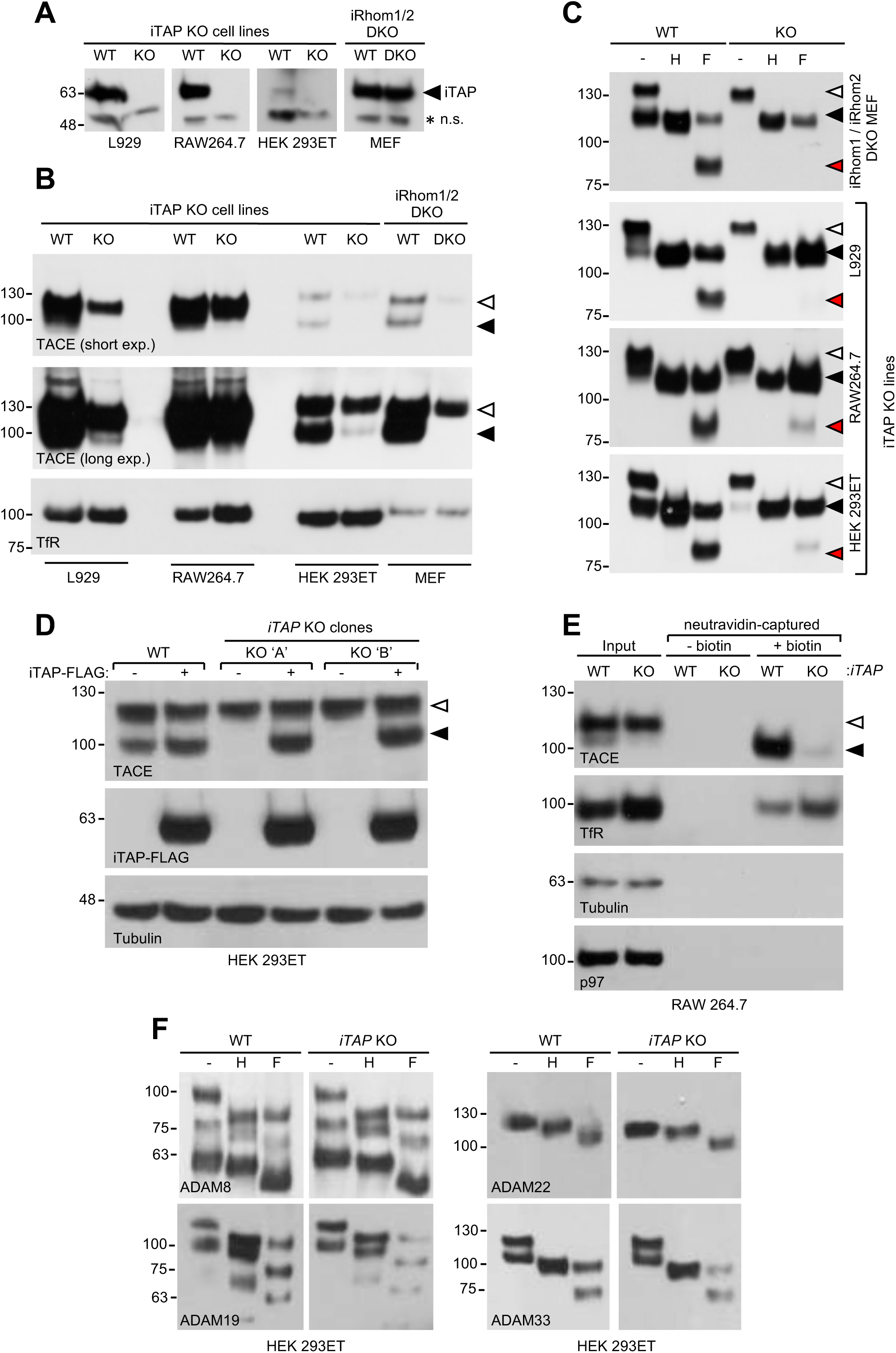
iTAP KO cells are specifically depleted in mature TACE levels. (**A**) iTAP was knocked out in L929, RAW 264.7 and HEK 293ET cells using CRISPR. Lysates were immunoblotted with anti-iTAP antibodies A non-specific band (n.s.) serves as a loading control. (**B**) Glycoproteins from the cells in (A) were enriched with using concanavalin A-sepharose (conA) and TACE levels were assessed by western blot. iRhom double KO MEFs were used as a reference and the transferrin receptor as a loading control. (**C**) Validation of mature and immature TACE detection in panels of WT versus iRhom2 DKO or iTAP KO cells, by deglycosylation. ConA enriched lysates from (A) were treated with endoglycosidase H (endo-H; H; which cleaves ER-resident glycans only) and PNGase F (F; which cleaves both ER and post-ER glycans). Here and throughout, the immature and mature species of TACE are indicated with white arrowheads and black arrowheads respectively, whereas red arrowheads denote the fully deglycosylated mature TACE polypeptide. (**D**) iTAP expression restores the presence of mature TACE in iTAP KO cells. Lysates from WT or iTAP KO HEK 293ET stably expressing empty vector or human iTAP were screened for TACE maturation. Tubulin was used as a loading control. (**E**) iTAP KO cells lack mature cell surface TACE. RAW 264.7 WT or iTAP KO were surface biotinylated *in vivo* and lysates were enriched for biotinylated proteins with neutravidin resin. The transferrin receptor was used as a cell surface positive control protein and tubulin to demonstrate that cytoplasmic proteins were not labelled. (**F**) iTAP is essential specifically for TACE maturation and not for the maturation of other ADAMs. HEK 293ET WT or KO cells were transfected with the indicated panel of V5-tagged ADAMs. The lysates were deglycosylated as described above.

### iTAP is required to maintain iRhom2 stability

The observation that iTAP-null cells exhibited drastically depleted mature TACE levels could be explained by two potential mechanisms. First, loss of iTAP, which binds to iRhom2 (Figures 1-2; Table S1), could impair ER exit of the iRhom/TACE complex, causing a failure in TACE maturation, as observed in iRhom KO cells. Alternatively, as TACE undergoes constitutive recycling from the plasma membrane (Lorenzen et al., 2016) and iRhom2 traffics to the cell surface and enters the endolysosomal pathway (Maney et al., 2015; Grieve et al., 2017; Cavadas et al., 2017), iTAP could stabilize iRhom/TACE complexes in the endocytic pathway or plasma membrane. Given the established role of FERM domain proteins on the cell cortex, this second possibility seemed plausible. To investigate the impact of iTAP on iRhom2 stability, we first used RAW 264.7 cells. These macrophage-like cells express high levels of endogenous iRhom2, making its detection more feasible than in HEK 293ET cells or MEFs. Strikingly, endogenous iRhom2 was depleted in iTAP KO cells (Fig. 6A) showing that iTAP is essential to maintain iRhom2 levels. Consistent with this, in HEK 293ET cells, iTAP transient overexpression increased the steady state levels of overexpressed iRhom2-HA and enhanced the stability of the protein, during a timecourse of cycloheximide treatment used to block additional protein synthesis (Fig. 6B). This experiment, in which iRhom2 was expressed from an artificial promoter, indicates that impact of iTAP on iRhom2 levels is independent of transcription. As predicted by these results, transiently overexpressed iRhom2-HA was destabilized in iTAP KO cells (Fig. S2A).

**Figure 6.**
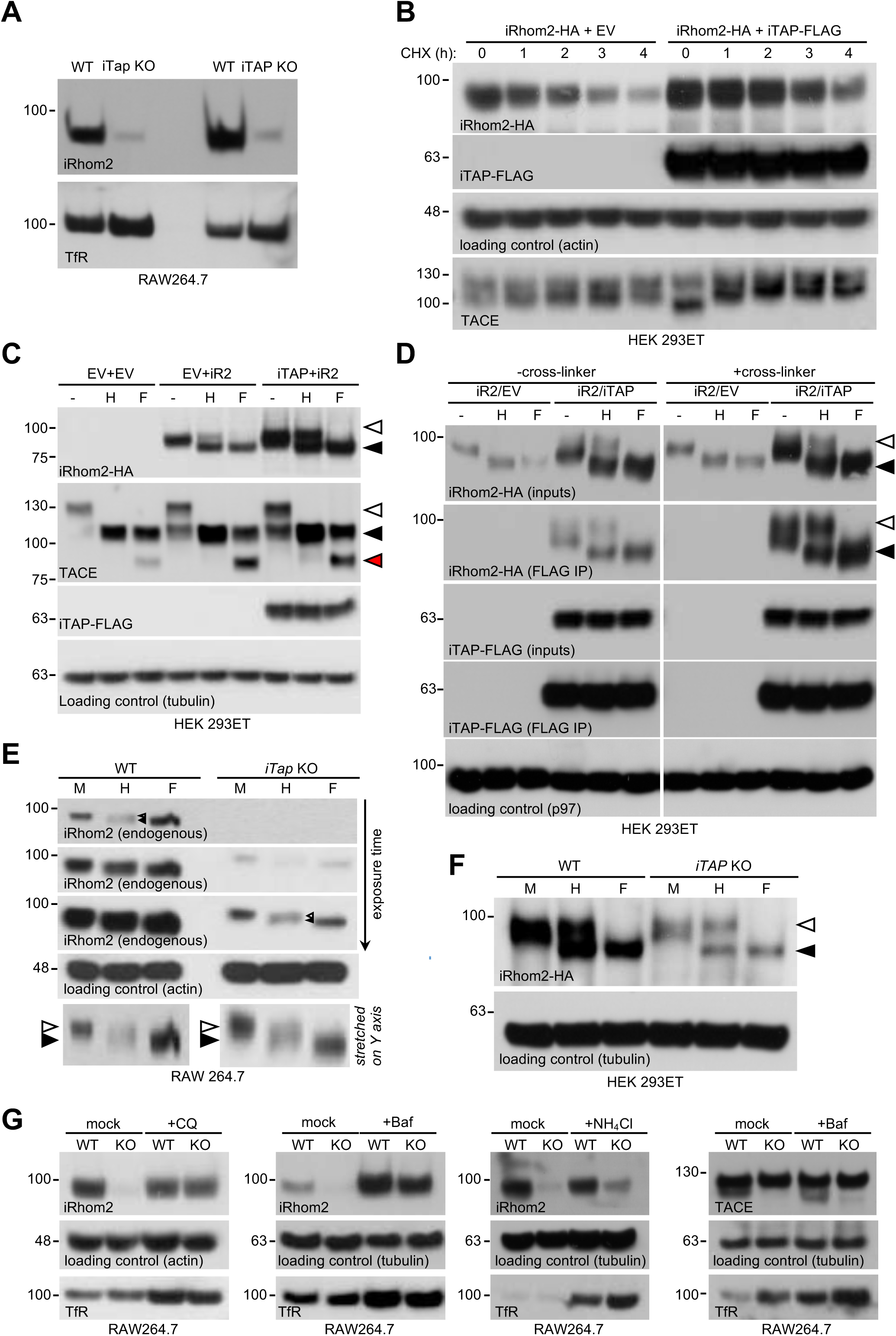
iTAP is required for iRhom stability preventing its degradation in lysosomes. (**A**). iRhom2 is depleted in iTAP KO cells. Lysates from WT vs iTAP KO RAW 264.7 were probed for endogenous iRhom2. The transferrin receptor (TfR) is a loading control. (**B**). iTAP expression enhances the stability of iRhom2. Stable iRhom2-HA-expressing HEK 293ET were transiently transfected with empty vector or iTAP-FLAG. 48h post transfection, the cells were treated with 100 μg/mL Cycloheximide (CHX) for the indicated durations. The stability of iRhom2 was assessed by HA blotting. (**C**). iTAP expression enhances the post-ER form of iRhom2. HEK 293ET expressing stably iRhom2-HA minus or plus iTAP-FLAG were deglycosylated as above. (**D**). The same cell lines were treated −/+ the cell-permeable crosslinker, DSP, and then anti-FLAG immunoprecipitations were performed from the lysates. The co-immunoprecipitated iRhom2-HA was deglycosylated as described before. (**E,F**). ER egress of iRhom2 is not impaired in iTAP KO cells. (**E**). Deglycosylation of endogenous iRhom2 in lysates from WT versus iTAP KO RAW 264.7 cells. The lysates were run on 4-12% gradient gels. The arrowheads in the endo-H lanes denote a doublet containing endo-H sensitive (upper arrowhead) and insensitive (lower arrowhead) species of iRhom2 detected in both WT and iTAP KO cells. The two panels at the bottom are cropped from the upper (WT) versus 3^rd^ from top (KO) iRhom2 exposures respectively. The two cropped images were grouped together, then artificially stretched, to the same degree, along the Y axis to accentuate the difference in iRhom2 mobility in response to endo-H versus PNGase F. (**F**). WT or iTAP KO HEK 293ET were transiently transfected with iRhom2-HA. Their lysates were deglycosylated as described. Endo-H–sensitive (black arrowhead) and -insensitive (white arrowhead) bands are noted. (**G**). iTAP prevents precocious sorting to the lysosomes. WT or iTAP KO RAW 264.7 cells were treated with 50 μM Chloroquine (CQ) for 48h, 100 μM Bafilomycin (Baf) for 16h, or 10 mM ammonium chloride (NH_4_Cl) for 48h and endogenous iRhom2 levels were detected by western blotting. Actin or tubulin were used as loading controls and the TfR acts as a control for the inhibition of lysosomal proteolysis. (**right panel**) Lysates from cells treated with Bafilomycin as described above, were conA-enriched and probed for TACE on a western blot. Tubulin and TfR are controls for loading and lysosomal inhibition, respectively. Throughout: Endo-H -sensitive (black arrowhead) and -insensitive (white arrowhead) bands are noted.

### iTAP promotes iRhom2 stability in the late secretory pathway

To address within which compartment iTAP acts on iRhom2 to govern its stability, we used deglycosylation with Endo-H, to discriminate between the pool of iRhom2 found in the ER, versus the Endo-H-insensitive fraction that has transited to the later secretory pathway. As anticipated (Zettl et al., 2011), the majority of overexpressed iRhom2 is found in the early secretory pathway, whereas iTAP expression, which increased the overall levels of iRhom2, substantially augmented the post-ER proportion (Fig. 6C).

To obtain insights into the compartment where iRhom2 and iTAP interact, and to provide hints about mechanism, we examined whether ER versus post-ER species of iRhom2 bound to iTAP. To discriminate between interaction *in vivo*, versus adventitious binding post-lysis, IPs were performed using lysates from cells treated with the cell-permeable cross-linker DSP, to covalently trap complexes of iTAP and iRhom2 *in situ* (Adrain et al., 2012). Notably, compared to IPs done without cross-linking, IPs from cross-linked cells showed a pronounced enrichment for the post-ER form of iRhom2 (Fig. 6D). Although we cannot exclude that iTAP binds to iRhom2 in the ER, iTAP shows a preference for the post-ER form of iRhom2, suggesting that its major role is to stabilize iRhom2 in the late secretory pathway

### iTAP is not required for iRhom2 egress from the ER but promotes iRhom2 stability on the cell surface

If iTAP stabilizes iRhom2 in the late secretory pathway, this predicts that iRhom2 should be able to transit from the ER in iTAP KO cells. We tested this hypothesis by assessing the partitioning within the secretory pathway of the traces of endogenous iRhom2 remaining in iTAP KO RAW 264.7 cells. (Fig. 6E). Glycosylated endogenous iRhom2 (Adrain et al., 2012) is more difficult to resolve electrophoretically than overexpressed tagged-iRhom2. Nonetheless, Endo-H digestion revealed an iRhom2 doublet comprising the ER-localized and post-ER forms, which could only be fully deglycosylated by digestion with PNGase F (Fig. 6E). We observed similar results with transfected iRhom2-HA in iTAP KO HEK cells (Fig. 6F). These data confirm that iTAP is not essential for the ER exit of iRhom2, and implies that it instead governs iRhom2 stability in the late secretory pathway. Consistent with this, when we performed live cell imaging experiments (as opposed to the experiments using fixed/permeabilized cells, shown in Fig. 3) we observed that the co-expression of iTAP-mCherry with eGFP-iRhom2 increased the amount of iRhom2 found on the cell surface (Fig. S2B). This demonstrates that iTAP increases the plasma membrane stability of iRhom2.

### iTAP modulates the stability of iRhom2 and TACE by preventing their precocious sorting to the lysosome

As iRhom2 leaves the ER in iTAP KO cells prior to its degradation, this predicts that loss of iTAP should trigger the degradation of iRhom2 and TACE in the late secretory pathway, i.e., in lysosomes. Consistent with this, a panel of lysosomotropic drugs that inhibit lysosomal proteolysis by impairing its acidification, rescued the iRhom2 stability in iTAP KO cells (Fig. 6F). In agreement with the premise that iTAP acts directly on iRhoms, the rescue of mature TACE under similar conditions was consistently more modest (Fig. 6F and data not shown), perhaps because of the slow trafficking time of TACE in the secretory pathway and because iTAP acts directly on iRhoms (Schlondorff et al., 2000). In conclusion, our data reveal that iRhom2 is mis-sorted into the lysosome, then degraded, in iTAP KO cells, highlighting an important physiological role for iTAP in maintaining the stability of iRhom2/TACE in the late secretory pathway.

### iTAP controls the stability of mature TACE, in vivo, in mice

iTap is expressed in a range of mouse tissues relevant to iRhom and TACE biology (Peschon et al., 1998, Li et al., 2015) (Fig. S1A). As the experiments conducted so far focused on transformed cell lines, we next examined the physiological importance of iTAP at the organismal level. To examine the role of iTap in mice, we generated a mutant in which the first coding exon of the *Frmd8* gene was deleted by CRISPR (Fig. 7A and Fig. S3). As anticipated, MEFs isolated from *iTap* KO embryos lacked iTap protein expression, confirming the successful targeting of the *iTap* gene (Fig. 7B). As anticipated, MEFs isolated from two independent *iTap* KO embryos lacked iTap protein expression (Fig. 7B) and exhibited a pronounced depletion in mature TACE levels (Fig. 7C).

**Figure 7.**
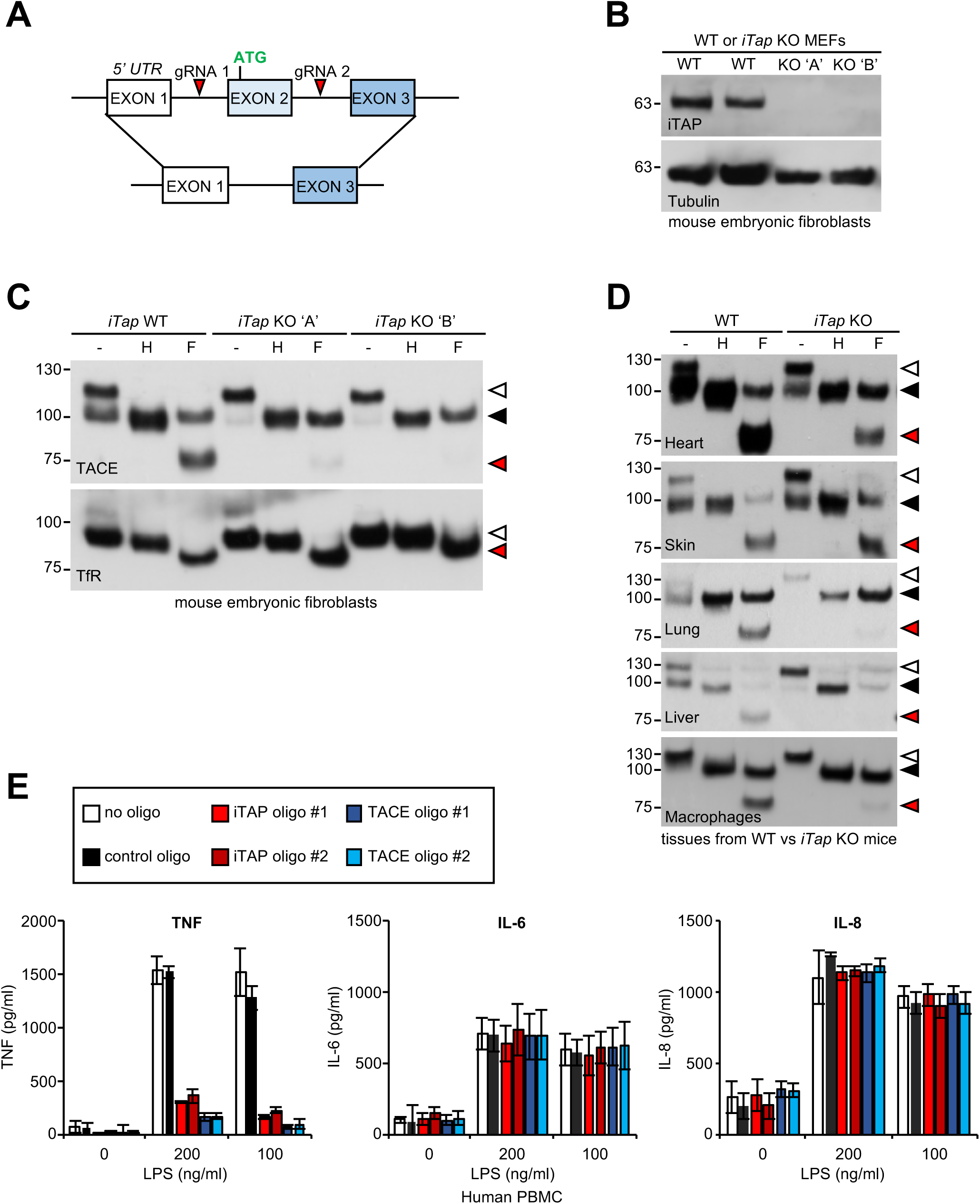
iTAP is essential for TACE maturation and function in primary cells and tissues from human and mouse. (**A**). Schematic representation of the CRISPR targeting strategy to delete mouse *Frmd8*/*iTap* gene using two guide RNAs flanking the first coding exon (exon 2). (**B**). Mouse embryonic fibroblasts were isolated from WT versus two independent *iTap* KO E14.5 embryo littermates. The loss of iTap at the protein level is shown by immunoblotting. (**C**). ConA-enriched lysates from WT versus *iTap* KO MEFs were deglycosylated as described previously. The transferrin receptor (TfR) is used as a loading control. (**D**). Mature TACE is depleted or diminished in TACE-relevant tissues from iTAP KO mice. ConA-enriched lysates from WT vs *iTap* KO mouse tissues and bone marrow-derived macrophages, were deglycosylated as described previously. TACE was detected by western blot. The immature and mature species of TACE are indicated with white arrowheads and black arrowheads respectively, whereas red arrowheads denote the fully deglycosylated mature polypeptide. The experiment was performed twice with lysates isolated from tissues from two individual KO mice. (**E**). iTAP is essential for TACE physiological regulation in human primary cells. Isolated primary human peripheral blood mononuclear cells (PBMC) were differentiated into monocytes, then electroporated with the indicated siRNAs. Cells were then stimulated with the indicated concentrations of lipopolysaccharide (LPS). After 18h, the concentration of the cytokines TNF, IL-6 and IL-8 secreted into the supernatants was measured by ELISA. The experiment was done three independent times and data from one representative experiment is shown. Data presented as mean ± standard error from triplicate measurements.

We next attempted to focus on potential phenotypes of the *iTap*–null mouse mutants themselves. As shown in Table S2, preliminary data from crosses between *iTap* heterozygous animals suggested that loss of iTap results in a degree of lethality during embryogenesis, resulting in sub-mendelian KO ratios when assessed at P1. Nonetheless, we successfully recovered a small number of live animals that escaped lethality. Notwithstanding the risk that these escapers may have engaged compensatory mechanisms to mitigate the loss of iTap, we harvested tissues to assess the maturation status of TACE (Fig. 7D). Significantly, with the possible exception of skin, we observed a substantial depletion in the relative proportion of mature TACE in a range of *iTap* KO mouse tissues, and in primary macrophages isolated from the bone marrow of iTap KOs. These data reinforce the notion that iTAP is an important physiological regulator of the iRhom/TACE/TNF axis *in vivo*, making it important to dissect fully, in future, the organismal role of iTAP.

### iTAP is a physiological regulator of TNF release in humans

As the PMA-stimulated release of a chimeric alkaline phosphatase-tagged TNF was impaired in iTAP KO cells (Fig. 4B), we hypothesized that iTAP was an important physiological regulator of TNF secretion. To test this, we derived primary monocytes from peripheral human blood, then differentiated them into macrophages. Notably, the stimulated release of endogenous TNF in response to lipopolysaccharide in these primary cells was profoundly impaired, when iTAP expression was ablated by siRNAs (Fig. 7E). As expected, secretion of IL-6 and IL-8, which is TACE-independent, was unaffected (Fig. 7E). Hence, iTAP is an essential physiological regulator of TNF secretion in human primary macrophages, the principal source of secreted TNF *in vivo*.

## Discussion

Our work identifies iTAP as an important physiological regulator of the iRhom2/TACE axis, which is essential for TNF secretion. As TACE is also the sheddase for a panoply of other substrates including ligands of the epidermal growth factor receptor, iTAP emerges as an important potential regulator of inflammation and growth factor signaling, during development, normal physiology, infection and inflammatory disease. We show that the binding of iTAP is essential to stabilize iRhom2 in the late secretory pathway, licensing it to promote the cell surface expression of mature TACE, driving signalling (summarized in Fig. S4).

Clearly, delineation of the physiological role of iTAP requires an extensive phenotypic analysis of *iTap* KO mice. Unfortunately, the partially penetrant lethality of *iTap* KO mice, and embryos, limits an immediate definitive analysis of the organismal phenotype, placing these experiments within the scope of future studies and potentially requiring the creation of a conditional *iTap* mutant mouse. In spite of these limitations, we show that mature TACE levels are dramatically depleted in tissues isolated from surviving *iTap* KO pups that escape lethality (Fig. 7D). This reinforces the notion that iTAP is an important physiological regulator of the iRhom/TACE/TNF axis *in vivo*, making it important to dissect fully, in future, the organismal role of iTAP.

An obvious question concerns to what extent the established features and roles of FERM domain proteins apply to iTAP and hence what can be predicted from this, about the regulation of the iRhom/TACE pathway? A general theme is that FERM domain proteins connect the cytoplasmic tails of cell surface client proteins to the cortical actin cytoskeleton, to enhance their stability (Hoover and Bryant, 2000; Baines et al., 2014; Moleirinho et al., 2013). While iTAP clearly binds to the cytoplasmic tails of iRhoms, which are found on the plasma membrane, our preliminary experiments failed to detect robust binding of iTAP to actin (Fig. S5A). Besides, we have not identified predicted actin binding motifs in the C-terminus of iTAP.

Future experiments will be required to determine precisely how iTAP stabilizes iRhom and TACE in the late secretory pathway. Notably, some FERM domain proteins are implicated in endosomal sorting, the process whereby endocytosed proteins are sorted in early endosomes, for routing to the multi-vesicular body, lysosome, *trans-*Golgi network, recycling endosome or alternatively, ‘fast’ recycling back to the cell surface (Cullen, 2008). Analogous to the degradation of iRhom2 in iTAP KO cells, loss of Snx17, which binds to the cytoplasmic tail of 1 integrins, results in a failure in their endocytic recycling, and lysosomal degradation (Böttcher et al., 2012). However, although sorting nexins are intimately connected with the endocytic recycling machinery, this is only one theme within the wider FERM domain family and it cannot be assumed that iTAP directly controls endocytic recycling. Future studies are required to clarify the relationship, between iTAP and the trafficking machinery, to map the vesicular itinerary taken by iRhom/TACE complexes, and to establish the basis of the mis-sorting defect in iTAP-null cells.

It will be interesting to reconcile the role of iTAP in the control of iRhom/TACE stability, versus that of PACS2, which binds directly to TACE (Dombernowsky et al., 2015). iTAP and PACS-2 both impact on TACE stability, but iTAP can presumably only influence TACE stability when the latter is bound to iRhoms. This is relevant because TACE stimulants trigger detachment of TACE from iRhom2 on the cell surface (Grieve et al., 2017), a mechanism important for facilitating access of TACE to its substrates (Cavadas et al., 2017). As iRhom and TACE are uncoupled at a crucial stage during signaling, their degradative fates could also be separated, leaving open the possibility that iTAP and PACS-2 may govern different stages in TACE's trafficking lifecycle.

The TNF (and EGFR) pathway(s) are very stringently regulated by positive and negative feedback (Avraham and Yarden, 2011; Wallach, 2016; Vereecke et al., 2009). Considering the significant impact that iTAP has on TACE, it is tempting to speculate that feedback into the signaling flux in the TNF release pathway could be governed by controlling the interaction between iTAP and iRhom, or even by modulating the stability of iTAP itself. Future studies are required to investigate this premise.

Inhibiting the activity of TACE has been the subject of considerable pharmaceutical interest for decades, but attempts have failed, often because of cytotoxicity associated with unintended collateral targeting of ADAMs and matrix metalloproteases, that share active site architectures related to TACE (Murumkar et al., 2010). As iTAP is a specific regulator of TACE pathways controlled by iRhoms, this makes the blockade of the iRhom:iTAP interaction an interesting potential future novel therapeutic approach for attenuating TACE activity and TNF secretion, during disease.

## Materials and Methods

### Reagents

The following reagents were used: 1,10-phenanthroline (131377, Sigma), Bafilomycin A1 (sc-201550, Santa Cruz), Chloroquine (C6628, Sigma), Ammonium chloride (Acros, 10676052), Ionomycin (Cayman, CAYM11932-5), Lipopolysaccharide E. coli 055:B5 (sc-221855A, Santa Cruz), Phorbol 12-myristate 13-acetate (PMA, P1585, Sigma), Cycloheximide (sc-3508, Santa Cruz), 1-Step PNPP Substrate (PIER37621, Thermo Fisher), Dithiobis (succinimidyl proprionate) (DSP, 10731945, Pierce), Sulfo-NHS-LC-Biotin (Thermo Scientific, 21335), NeutrAvidin Protein (11885835, Thermo Fisher), Concanavalin A Agarose (786-216, G-biosciences), Endo-Hf (174P0703, NEB), PNGase F (174P0704, NEB), TOPO TA Cloning Kit for Sequencing, (450030, Invitrogen), Gibson Assembly Master Mix (174E2611, New England Biolabs), KOD Hotstart DNA polymerase (71086-5, Novagen). Polyethylenimine branched (408727, Sigma), FuGENE 6 (2691, Promega), Opti-MEM I Reduced Serum Medium, GlutaMAX (51985-026, Life technologies), NuPAGE Novex 4-12% Bis-Tris Protein Gels 1.0 mm (Novex, NP0322BOX, Life Technologies).

### Antibodies

The following antibodies were used: TACE (Ab39162, Abcam); TACE Ab318 (monoclonal anti-TACE antibody (Trad et al., 2011) was used for the detection of mouse TACE; alpha-Tubulin (sc-8035, Santa Cruz), alpha-Tubulin (Clone YL1/2, Ana Regaldo, IGC), p97 (Thermo, MA1-21412), HA-HRP (clone 3F10, Roche), HA (clone HA.11, 901501, Biolegend) V5-HRP (R961-25, Life Technologies), Frmd8 (157H00083786-B01P, Abnova), Transferrin Receptor (13-6800, Life Technologies), Flag-HRP (A8592, Sigma), GAPDH (2118, CST), Actin (abcam ab8227). Anti-iRhom2 polyclonal antibodies specific to the mouse iRhom2 N-terminus (amino acids 1-373) or raised against the iRhom homology domain, were previously described (Adrain et al., 2012). For immuno-precipitations, magnetic beads were used: anti-HA (15222405, Pierce), ANTI-FLAG^®^ M2 Affinity Gel (A220, Sigma), MagnaBind Goat Anti-Mouse IgG Beads (21354, Thermo Scientific), MagnaBind Goat Anti-Rabbit IgG Beads (21356, Thermo Scientific).

### Plasmids

C-terminally triple HA-tagged versions of human iRhom1, the cytoplasmic N-terminus of iRhom1 (amino acids 1-404), mouse iRhom2, mouse RHBDD2, human RHBDD3 and mouse UBAC2 were cloned into the lentiviral expression plasmid pLEX-MCS, using Gibson cloning. These plasmids were used only for the respective mass spectrometry experiments. HA-tagged mouse iRhom2, iRhom2 ∆IRHD, human iRhom1, the cytoplasmic N-terminus of iRhom1, mouse Ubac2 and mouse Rhbdd2 were cloned into the retroviral vector pM6P (a kind gift of Felix Randow). C-terminally triple FLAG-tagged iTAP was cloned into pM6P vector using Gibson assembly. The N-terminal truncations of iRhom2 used in Figure 2 were cloned into a modified version of the lentiviral expression vector pLEX-MCS in which the puromycin resistance cassette was replaced with a blasticidin resistance gene. The packaging vectors for the production of retrovirus or lentivirus were described previously (Cavadas et al., 2017). The mCherry-tagged iRhom2 plasmid was previously described (described in Luo et al. 2016 and human iTAP-GFP was inserted with standard cloning techniques into pIC111 vector (a gift from Iain Cheeseman & Arshad Desai; Addgene plasmid # 44435). Alkaline phosphatase-tagged TACE substrates, a gift of Shigeki Higashiyama were described previously (Sahin et al., 2004). V5-tagged ADAM expression plasmids and secreted luciferase construct were described previously (Christova et al., 2013). CRISPR plasmids are described below. Human Tumor Necrosis Factor (TNF) containing an N-terminal FLAG tag, was cloned into pCR3 by standard techniques. Flag-tagged SREBP2 was a gift of Larry Gerace and STING-FLAG a gift of Lei Jing). In supplemental figure 4B, iRhom2 in a modified version of pEGFP-N1 was used as a template for Quick-Change mutagenesis resulting in the constructs iRhom2 NPAY>AAAA, iRhom2 NRSY>AAAA and iRhom2 Double NxxY>AAAA.

### Cell culture

HEK 293ET, RAW 264.7, L929 and MEFs were maintained under standard conditions in Dulbecco's Modified Eagle Medium (DMEM)-high glucose supplemented with fetal bovine serum. Bone Marrow Derived Macrophages (BMDM) were isolated from 8-week old mice and cultured as previously described (Adrain et al., 2012). Embryonic fibroblasts were generated from E14.5 embryos and immortalized using lentiviral transduction of SV40 virus large T antigen.

### Cytokine secretion in isolated human monocytes

Primary human peripheral blood mononuclear cells (PBMC) were purified from donor whole blood using the Ficoll-Hypaque gradient method as described previously (Henry et al., 2016). After overnight plastic adherence in heat-inactivated serum containing medium, non-adherent cells were removed and remaining cells were washed three times in PBS. Macrophage differentiation was induced using recombinant human macrophage-colony stimulating factor (M-CSF, 100 ng/ml) over five-seven days during cell culture in RPMI supplemented with 10% FCS. Primary human macrophages (5 × 10^5^) were nucleofected with 200 nM of each siRNA (control NS oligo, MWG Eurofins − 5′-GUUCCUGAGCCUGGACUAC −3′; iTAP oligo #1, Santa Cruz - catalog code sc-96500; iTAP oligo #2, GE Dharmacon - M-018955-01-0005; TACE oligo #1, Santa Cruz - sc-36604; TACE oligo #2, GE Dharmacon - M-003453-01-0005) in nucleofection buffer (5 mM KCl, 15 mM MgCl_2_, 20 mM HEPES, 150 mM Na_2_HPO_4_ [pH 7.2]) using Amaxa Nucleofector (program Y-010). Cells were plated in 6-well plates (2 ×10^5^ cells/well) or in 24-well plates (1 × 10^5^ cells/well) and 48 h after nucleofection were stimulated with lipopolysaccharide (LPS). After 18 h, cell culture supernatants were collected and clarified by centrifugation for 5 min at 800 × g. Cytokines and chemokine concentrations were measured from clarified cell culture supernatants using specific ELISA kits obtained from R&D Biotechne systems (human TNF – DY210; human IL-6 – DY206; IL-8 – DY208).

### Retroviral transduction

HEK 293ET cells (1 × 10^6^) were transfected with pCL (−Eco, or 10A1) packaging plasmids (Naviaux et al., 1996) plus pM6P.BLAST empty vector (kind gift of F. Randow, Cambridge, UK) or pM6P containing the cDNA of human or mouse iTAP or mouse iRhom2. WT or iTAP KO HEK 293ET cells were transduced with the viral supernatant supplemented with polybrene 8 μg/mL, and selected with blasticidin (8 μg/mL) to generate stable cell lines. To transduce RAW 264.7 or L929, lentiviruses were prepared and concentrated as follows: HEK 293FT cells (24 × 10^6^) were transfected with pMD-VSVG envelope plasmid, psPAX2 helper plasmid and pLentiCas9-blast or empty vector pLentiGuide-puro or pLentiGuide-puro containing iTAP targeted gDNAs. The viral supernatants were concentrated 300-fold using ultracentrifugation (90,000 g) at 4°C for 4h, followed by re-suspension in 0.1 % BSA in PBS. Cells were transduced with the concentrated virus, supplemented with 8 μg/ml polybrene.

### Generation of iTAP KO cells via CRISPR

For CRISPR-mediated knockout of iTAP in human cells, gRNAs targeting exons common to all transcripts: the 1^st^ coding exon 5’-GCCCCGCTGAGCGATCCCAC-3’ or coding exon 4 of iTAP 5’-ACGTGTTCTTCCCAAAGCGG-3’ were cloned into plentiCRISPR v2 (Addgene plasmid # 52961), a gift from Feng Zhang. For the ablation of iTAP in human cells, HEK 293ET cells (2.5 x 10^4^ / cm^2^) were transfected using Fugene with pLentiCRISPR v2 empty vector, or either of the pLentiCRISPR-derived sgRNA plasmids described above. The next day, the cells were selected with puromycin (8μg/mL) for 3 days until mock transfected cells were eliminated. Cells were expanded and single-cell sorted by FACS or serial dilutions on 10 cm culture plates. To screen for the presence of indels in clones, genomic DNA was extracted from each clone and a 200 bp region flanking the site targeted by the gRNA was amplified for exon 1 (forward = CCTCCAGCCCCCCATCCCTGGCTC; reverse = GCCAGAGCTACTTCTCCAGGGCTGGGG) or exon 4 (forward = TCGGGAGAGGGGAGGGCTAAGCAG; reverse = GGGCAAGGTGCGAATGTCCAGGGGTC). Clones with mutant alleles were selected and the original PCR fragments amplified were isolated and sequenced via TOPO TA cloning. The selected clones “KO A” and “KO B” which contain indels in all alleles of iTAP, were then confirmed for loss of iTAP at the protein level by immunoprecipitation and subsequent western blot with an anti-FRMD8 antibody.

To ablate iTAP in mouse cells, gRNAs targeting the first coding exon (TTCGGTGGGACCGCTCCGCA) or second coding exon (GCACTACTGTATCATCCGCC) were cloned into plentiGuide-Puro (Addgene plasmid # 52963) and used in combination with lentiCas9-Blast (Addgene plasmid # 52962); both gifts of Feng Zhang. For transfection of the gRNAs, 2 × 10^5^ L929 or 5 x 10^5^ RAW 264.7 cells were transduced with 40 μl (RAW 264.7) or 20 μl (L929) of 300-fold concentrated lentivirus encoding pLentiCas9-Blast (Addgene #52962) and selected with blasticidin (4 μg/mL, RAW 264.7 and 8 μg/mL, L929). The Cas9-expressing lines were then transduced with the pLentiGuide-Puro sgRNA plasmds targeting the 1^st^ or 2^nd^ coding exons of mouse iTAP. Following selection with puromycin (4 μg/mL, RAW 264.7 and 7 μg/mL L929) the cells were single clone sorted by FACS. To screen for iTAP KO clones, genomic DNA was extracted from each clone and PCR used to amplify a 200bp region flanking the guide sequence (exon 1: forward=TTGAGAGCTTGAGGAGACCA; reverse CAGGCTGGAACCAAAGAGTTC); exon 2: forward=GGAAATGCTGATTGGACCTC; reverse CCTGCTGCCAGACCTTACCC). Clones with mutant alleles were identified as described for human cells.

### Experiments with mice

Experiments with mice were performed in accordance with protocols approved by the Ethics Committee of the Instituto Gulbenkian de Ciência and the Portuguese National Entity (DGAV-Direção Geral de Alimentação e Veterinária) and with the Portuguese (Decreto-Lei no. 113/2013) and European (directive 2010/63/EU) legislation related to housing, husbandry, and animal welfare.

### Generation of *iTAP* mutant mice

iTAP mutant mice were generated via CRISPR/Cas9 as previously described (Wang et al., 2013; Casaca et al., 2016). In brief, two gRNA´s (5’-CAGCCGAGTGCAGATCGGGT-3’ and 5’-GTGGCGGACTCAGAAATCAA-3’) were designed to introduce a deletion of the first coding exon (exon 2) of the mouse *Frmd8* gene. Oligos encoding the gRNA were inserted into the plasmid pgRNAbasic (Casaca et al., 2016), which contains a T7 promoter. The linearized vector was used as template for the production of sgRNAs, produced by *in vitro* transcription using the MEGAshortscript T7 Kit (Life Technologies). RNA was cleaned using the MEGAclear kit (AM1908, Life Technologies). Cas9 mRNA was produced by *in vitro* transcription using the mMESSAGE mMACHINE T7 Ultra Kit (Life Technologies) and plasmid pT7-Cas9 as a template (Casaca et al., 2016). Cas9 mRNA (10 ng/ml), plus the sgRNAs (10 ng/ml) were injected into the pronuclei of fertilized C57 BL/6 oocytes using standard procedures (Hogan et al., 1986). Deletions were assessed by PCR from tail genomic DNA using primers 5′-CCCGACTTGTTTGGCCATTTC-3′ or 5’-CGGGGCCTCGGGTTTG-3’ (forward) and 5′-TGGGACAAAGGAAGTGGTGCC-3′ (reverse). The deletion was confirmed by direct sequencing and TOPO cloning sequencing. These primers (along with 5’-ACTTTCACCCTACACATTTG-3’ 5’-AGTCCGCCACATCTAAAC-3’ for better amplification of WT alleles) were also used for genotyping mice and embryos of the *iTap* KO line.

### Immunostaining and fluorescence microscopy

HeLa (5 × 10^4^ cells/well) were plated on coverslips and transfected with iTAP-GFP (600 ng) or iRhom2-Cherry (600 ng), either alone or in combination. After 24 h, cell supernatant was removed and cells were washed 3 times with PBS (2 mL). Cells were fixed with 3 % paraformaldehyde for 10 minutes. Cells were washed again 3 times with PBS (2 mL) followed by permeabilisation with 0.15 % TX100 for 15 minutes. Cells were blocked with 2 % BSA (in PBS, pH 7.2) for 1 h to reduce non-specific binding of antibodies. Specific primary antibodies against Calnexin (Cell Signaling, C5C9) and Golgi GM130 (BD, 610823) were diluted 1:100 in 2 % BSA. Primary antibodies were incubated for 2 h at room temperature. Cells were washed 3 times with PBS (2 mL). Cells were incubated with the relevant rhodamine red-conjugated secondary antibody (Alexa Fluor) diluted 1:1000 in 2 % BSA for 1 h at room temperature. Cells were washed again with PBS, followed by incubation with Hoechst (Sigma) for 10 minutes. Coverslips were mounted on slides with 5 μL of Slow Fade (Molecular Probes).

For mitotracker staining, cells were transfected with iTAP-GFP (600 ng), as described previously. After 24 h, cells were treated with Mitotracker-Red (50 nM) for 15 minutes at 37 °, followed by fixation with 3 % paraformaldehyde. Nuclei were stained with Hoechst. Cells were visualised and analysed using a laser scanning confocal microscope (Olympus FV1000) using a 488 nm Argon laser (green fluorescence), a 543 nm HeNe laser (red fluorescence) and a 405 nm LD laser. Confocal images were acquired using Fluroview 1000 V.1 software.

### Live fluorescence microscopy

iRhom1/iRhom2 DKO MEFs stably expressing eGFP-mouse iRhom2 either alone or together with mouse iTAP-mCherry were plated (5 × 10^4^ per well) on 4-chamber glass-bottomed dishes (In Vitro Scientific, D35C4-20-1.5-N) 24 hours prior to imaging. Cells were imaged on a laser scanning confocal microscope Zeiss LSM 780* using the 40×/1.2 M27 W Korr C-Apochromat objective and a 488nm or 561nm excitation wavelength.

### *In vivo* protein cross-linking with Dithiobis Succinimidyl Proprionate (DSP)

Cells are washed twice in cold PBS before incubation in 0.2 mg/mL DSP for 45min. The cross-linker was aspirated off and the cell monolayers were washed three times for 10 min in ice cold PBS containing 50 mM Tris, pH 8.0 to quench any remaining cross-linker. Subsequently the cells were lysed in Triton X-100 lysis buffer (150 mM NaCl, 50 mM Tris-HCl, protease inhibitors, pH 7.4 and 10 mM 1,10-phenanthroline). Post nuclear supernatants were supplemented to contain 0.1% SDS and 0.25% sodium deoxycholate.

### Co-immunoprecipitations

HEK 293ET cells expressing the indicated plasmids were lysed for 10 minutes on ice in TX-100 lysis buffer (1% Triton X-100, 150 mM NaCl, 50 mM Tris-HCl, pH 7.4) containing complete protease inhibitor cocktail (Roche), and 10 mM 1,10-phenanthroline (to inhibit TACE autoproteolysis). Post-nuclear supernatants were pre-cleared with unconjugated magnetic beads or agarose at 4°C for 60 minutes with rotation, followed by capture on anti-HA magnetic beads or anti-Flag respectively for 90 minutes. Beads were washed 3-5 times, for 10 minutes, at 4ºC in the same Triton X-100 lysis buffer supplemented with NaCl to 300 mM. Samples were eluted with 1.5 x SDS-PAGE sample buffer and incubated at 65ºC for 15 minutes before loading.

### Identification of Rhomboid-interacting proteins

HEK 293ET cells were stably transduced with lentiviruses encoding pLEX empty vector, or pLEX derivatives containing HA-tagged iRhom1, iRhom2, iRhom1 N terminus, RHBDD2, RHBDD3, UBAC2. Live cells were washed twice with ice-cold PBS, then left untreated or treated with the crosslinker DSP (0.2 mg/mL) as described below. Lysates were clarified, then pre-cleared with irrelevant control antibodies conjugated to magnetic beads for 60 min at 4°C with rotation. After saving ‘input’ samples, the lysates were incubated with anti-HA resin for 90 min at 4°C with rotation. Subsequently, the precipitated beads were washed four times in the respective buffers indicated above. One quarter of the beads were reserved for SDS-PAGE analysis, whereas three-quarters of the precipitated beads were resuspended in UREA buffer (8M Urea, 4% CHAPS, 100 mM DTT, 0.05% SDS). For MS analysis, immunoprecipitates were enzymatically digested on 3 kD MWCO filters (Pall Austria Filter GmbH) using an adaption of the FASP protocol as described previously (Bileck et al., 2014; Slany et al., 2016). After pre-concentration of the samples, protein reduction and alkylation was performed, then trypsin was added and incubated at 37°C for 18h. The digested peptide samples were dried and stored at −20°C then later reconstituted in 5 μl 30% formic acid (FA) containing 10 fmol each of 4 synthetic standard peptides and further dilution with 40 μl mobile phase A (98% H_2_O, 2% ACN, 0.1% FA), as described previously (Bileck et al., 2014; Wiśniewski et al., 2009). LC-MS/MS analyses were performed using a Dionex Ultimate 3000 nano LC-system coupled to a QExactive orbitrap mass spectrometer equipped with a nanospray ion source (Thermo Fisher Scientific). For LC-MS/MS analysis, 5μl of the peptide solution were loaded and pre-concentrated on a 2 cm x75 μm C18 Pepmap100 pre-column (Thermo Fisher Scientific) at a flow rate of 10 μl/min using mobile phase A. Following this pre-concentration, peptides were eluted from the pre-column to a 50cm x75μm Pepmap100 analytical column (Thermo Fisher Scientific) at a flow rate of 300 nl/min and further separation was achieved using a gradient from 7% to 40% mobile phase B (80% ACN, 20% H_2_O, 0.1% FA) over 85min including column washing and equilibration steps. For mass spectrometric analyses, MS scans were accomplished in the range from m/z 400-1400 at a resolution of 70000 (at m/z =200). Subsequently, data-dependent MS/MS scans of the 8 most abundant ions were performed using HCD fragmentation at 30% normalized collision energy and analyzed in the orbitrap at a resolution of 17500 (at m/z =200). Protein identification was achieved using Proteome Discoverer 1.4 (Thermo Fisher Scientific, Austria) running Mascot 2.5 (Matrix Science). Therefore, raw data were searched against the human proteome in the SwissProt Database (version 11/2015 with 20.193 entries) with a mass tolerance of 50 ppm at the MS1 level and 100 mmu at the MS2 level, allowing for up to two missed cleavages per peptide. Further search criteria included carbamidomethylation as fixed peptide modification and methionine oxidation as well as protein N-terminal acetylation as variable modifications.

### Cell Surface Biotinylation

Biotinylation was performed as previously described for BMDM with small modifications (Adrain et al., 2012). RAW 264.7 macrophages (1.5×10^6^ cells, 6 well plates) were moved to a cold room (at 4°C), washed with ice-cold PBS pH 8.0 for 10 minutes, incubated with (1 mg/mL) Sulfo-NHS-LC-Biotin in PBS pH 8.0, according to the manufacturer's instructions. Following quenching with 50 mM Tris in PBS, cells were lysed for 10 minutes with TX-100 lysis buffer (1,10-phenathroline, protease inhibitors, 50 mM Tris), then biotin labelled surface proteins from post-nuclear supernatants were captured on neutravidin agarose resin at 4°C overnight. The resin was washed 3 times, 10 minutes, with TX-100 lysis buffer containing 300 mM NaCl at 4°C. Samples were eluted with 1.5x SDS-PAGE sample buffer and incubated at 65ºC for 15 minutes, before loading.

### Glycoprotein enrichment using Concanavalin A

To improve the detection of TACE, MEFs were lysed in TX-100 lysis buffer supplemented with 1 mM EDTA, 1 mM MnCl_2_, 1 mM CaCl_2_ and glycoproteins were captured using Concanavalin A (ConA) Agarose. Beads were washed twice in the same buffer and eluted by heating for 15 min at 65°C in sample buffer supplemented with 15% sucrose or for 5 min at 95°C in sample buffer supplemented with 15% sucrose and 1x Glycoprotein denaturation buffer (NEB).

### *In vivo* Protein Deglycosylation Analysis

Post nuclear lysate supernatants or denatured lysates, are denatured at 65°C for 15min in the presence of 1× Glycoprotein Denaturing buffer (NEB). Endo-H and PNGase F reactions are set up according to manufacturer's instructions for 1h at 37°C.

### Shedding and secretion assays

Shedding assays were performed using previously described plasmids encoding alkaline phosphatase-tagged EGFR ligands: Transforming Growth Factor α (TGFα), Amphiregulin (AREG), Epiregulin (EPIREG), Heparin Binding-Epidermal Growth Factor (HB-EGF), Epidermal Growth Factor (EGF) and Betacellulin (BTC) or Tumor necrosis factor (TNF) (Sahin et al., 2006; Zheng et al., 2002). HEK 293ET or MEF (3 × 10^5^ in 6 well-plates) were transfected with 1 μg cDNA of AP-substates and 6 μl PEI or 3 μL respectively. The next day cells were washed 3 times with serum free media before incubation for 1 hour in 1 mL Optimem (containing the vehicle of the drug in next step) (for basal shedding), followed by 1 hour with 1 mL Optimem containing 1 μM PMA or 2.5 μM ionomycin (for stimulated shedding). Supernatants from each incubation step were collected. Following, the cells were washed in ice-cold PBS three times and lysed in Triton X-100 buffer described previously (unshed material). Supernatants and lysates were centrifuged on a bench-top centrifuge at top speed for 10 min to remove cells and cell debris. Supernatants and lysates were incubated with the Alkaline Phosphate (AP) substrate p-nitrophenyl phosphate (pNPP) at room temperature. AP activity measured using a 96-well plate spectrophotometer (405 nm). Results are presented as PMA or IO “shedding over total” calculated by the formula ('stimulated shedding’-'basal shedding’) / ('stimulated shedding’+’unshed’). The release into the medium of a secreted form of luciferase containing a signal peptide was assessed as a control, as described (Christova et al., 2013).

### Protein Extraction from mouse tissues

Protein was extracted from mouse tissues by lysing in a modified RIPA buffer (150 mM NaCl, 50 mM Tris-HCl, pH 7.4, 1 mM EDTA, 1% Triton X-100, 1% Na Deoxycholic Acid, 0.1% SDS containing protease inhibitors and 10 mM 1,10-phenanthroline). Homogenates were clarified and normalized, then incubated with ConA resin, as described above.

### Statistical analysis

All statistical analyses were done with two-sample two tail unpaired t tests assuming unequal variances, using excel software. The comparisons were performed after transforming the raw values into their relative (fold change) values to the WT sample.

## Acknowledgements

We thank Felix Randow, Shigeki Higashiyama and Feng Zhang for plasmids. We thank Florian Steinberg for discussions and disclosure of unpublished results. We thank Matthew Freeman for helpful discussions. We express our deep gratitude to Moises Mallo for advice concerning CRISPR plus CRISPR reagents. We are grateful for the assistance of Ana Nóvoa and IGC's transgenics and mouse facilities. We thank IGC's cell sorting/flow cytometry, sequencing, and histopathology facilities. CA acknowledges the support of Fundação Calouste Gulbenkian, Worldwide Cancer Research (14-1289), a Marie Curie Career Integration Grant (project no. 618769), Fundação para a Ciência e Tecnologica (FCT, SFRH/BCC/52507/2014; PTDC/BEX-BCM/3015/2014), the European Crohn's and Colitis organization (ECCO), and COST BM1406. SJM acknowledges the support of Science Foundation Ireland (14/IA/2622). KS acknowledges the support of EMBO (Installation Grant no. 2329), Ministry of Education, Youth and Sports of the Czech Republic (project no. LO1302) and European Regional Development Fund (project OPPK no. CZ.2.16/3.1.00/24016). IO acknowledges the support of Fundação Calouste Gulbenkian – IGC (fellowship contract as ref. 91/BD/14). M.C. acknowledges the support of the FCT (grant SFRH/ BPD/117216/2016).

---

**Table S1.** iTAP interacts with endogenous iRhoms. Lysates from HEK 293ET cells expressing iTAP-FLAG versus cells containing empty vector or expressing a panel of control proteins (TNF-FLAG, STING-FLAG, SREBP2-FLAG) were immunoprecipitated for FLAG and subjected to mass spectrometry. Peptides assigned to iRhom-1 or iRhom2 are found specifically in iTAP precipitates.

| ***iRhom1*** | **Vector** | **iTAP** | **TNF** | **STING** | **SREBP2** |
| --- | --- | --- | --- | --- | --- |
| Peptide counts | 0 | **27** | 0 | 0 | 0 |
| Sequence coverage [%] | 0 | **37.7** | 0 | 0 | 0 |

| ***iRhom2*** | **Vector** | **iTAP** | **TNF** | **STING** | **SREBP2** |
| --- | --- | --- | --- | --- | --- |
| Peptide counts | 3 | **44** | 2 | 2 | 2 |
| Sequence coverage [%] | 2.9 | **45.4** | 1.8 | 2.2 | 2 |

**Table S2.** Mendelian ratios of embryos isolated at embryonic day 14.5 (E14.5), or pups at (P1) post-partum obtained from crosses obtained between *iTap* heterozygous mice.

| Developmental Stage | *iTap* +/+ | *iTap* +/- | *iTap* -/- | Total No. of Animals |
| --- | --- | --- | --- | --- |
| **E14.5** | 10 | 17 | 4 | 31 |
| **%** | 32,2 | 54,8 | 12,9 |  |
| **Ratio** | 1 | 1,7 | 0,4 |  |
| **P1** | 17 | 46 | 4 | 70 |
| **%** | 25,37 | 68,66 | 5,97 |  |
| **Ratio** | 1 | 2,71 | 0,24 |  |

**Figure S1.**
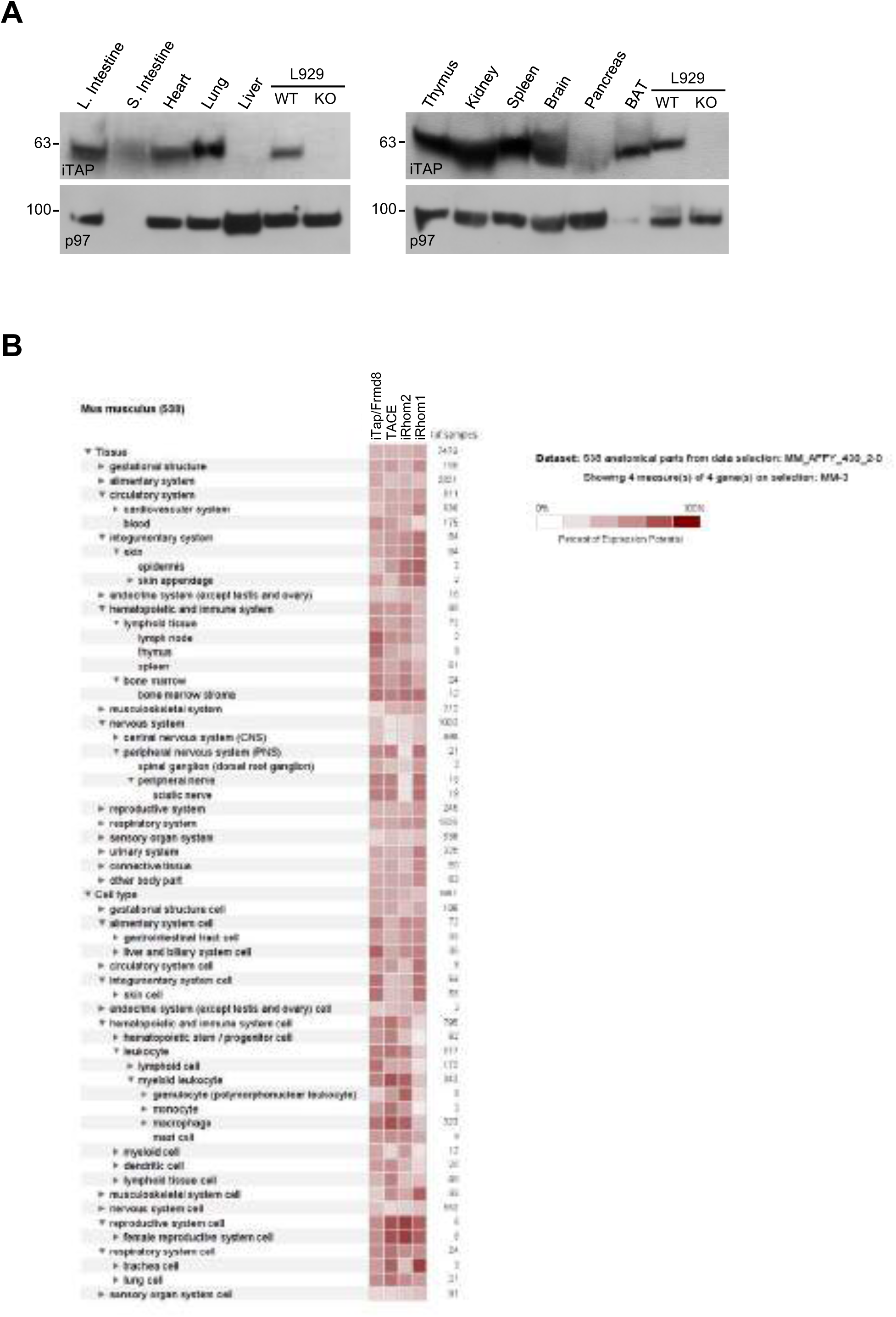
iTap is broadly expressed in a variety of tissues important for TACE biology. (**A**) Lysates were prepared from a panel of tissues from WT C57 BL/6J mice and iTAP was detected by western blot. Lysates from WT versus *iTap* KO L929 mouse fibroblasts were used as a control. (**B**) The extent of co-expression between iTap/Frmd8, TACE/ADAM17, iRhom2 or iRhom1 in mouse anatomical parts and cell types was investigated by interrogating gene expression data from the GeneChip™ Mouse Genome 430 2.0 Array via the Genevestigator platform (Hruz et al., 2008).

**Figure S2.**
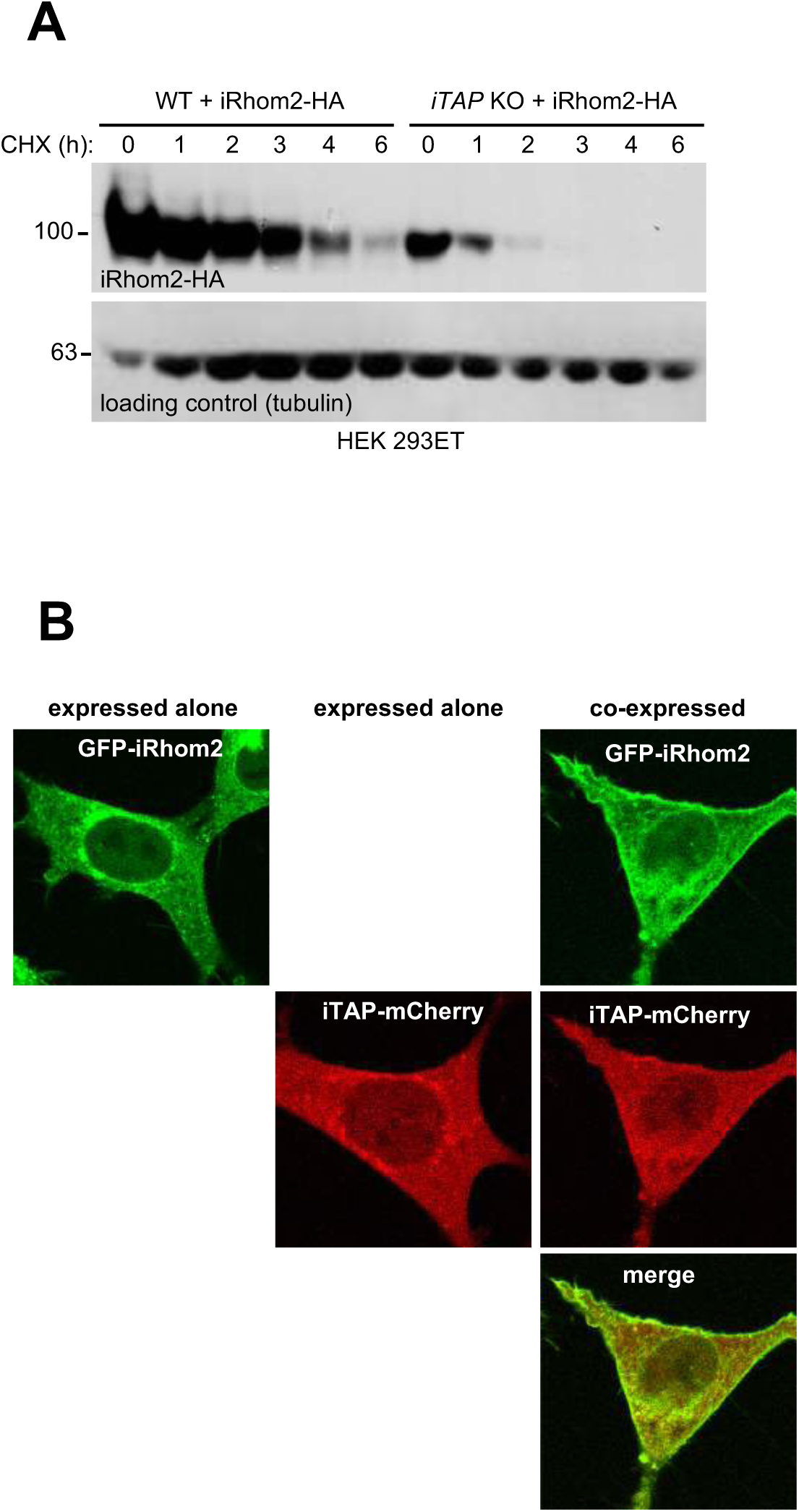
iTAP ablation increases iRhom2 degradation in the lysosomal compartment. **(A)** Absence of iTAP destabilizes iRhom2. WT or *iTAP* KO HEK 293ET cells were transiently transfected with iRhom2-HA. 48h post transfection, the cells were treated with 100 μg/mL CHX for the indicated durations. The stability of iRhom2 was then assessed by HA blotting. Tubulin was used as loading control. (**B**) iRhom1/2 DKO MEFs stably expressing eGFP-mouse iRhom2 either alone or together with mouse iTAP-mCherry on 4-chamber glass-bottomed dishes were imaged, as live cells, on a laser scanning confocal microscope (Zeiss LSM 780) using the 40×/1.2 M27 W Korr C-Apochromat objective and a 488 nm or 561 nm excitation wavelengths.

**Figure S3.**
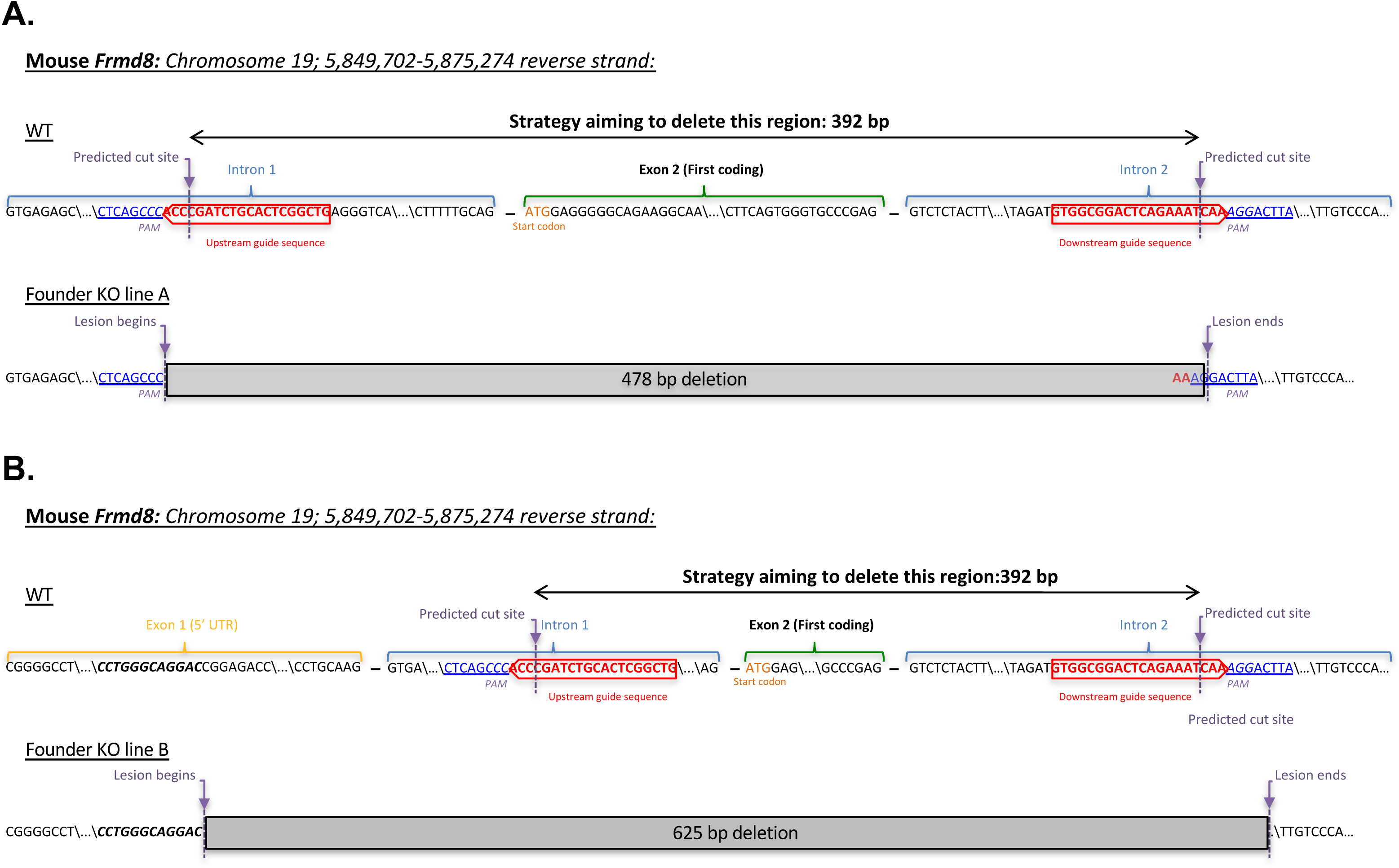
Mouse *Frmd8*/*iTap* gene targeting via CRISPR. A pair of gRNAs (indicated in red) was selected to engineer the deletion of the first coding exon of mouse *Frmd8/iTap* as described in materials and methods. The PAM sequence and the theoretical Cas9 cut sites 3-4 bp from the PAM are indicated. The resulting founder animals were genotyped from tail biopsies and the identity of the lesions revealed by DNA sequencing. Two lines of animals (A and B) were established following germline transmission of the mutant alleles from individual founders. KO line **A** contains a 478 bp deletion that removes exon 2 along with parts of introns 1 and 2. KO line **B** contains a larger deletion of 625 bp that starts within the non-coding exon 1, deletes all of intron 1, exon 2 and part of intron 2, as shown. In both cases, the precise nucleotide sequence on the 5’ and 3’ boundaries of the mutant lines is indicated and aligned with the WT genomic sequence.

**Figure S4.**
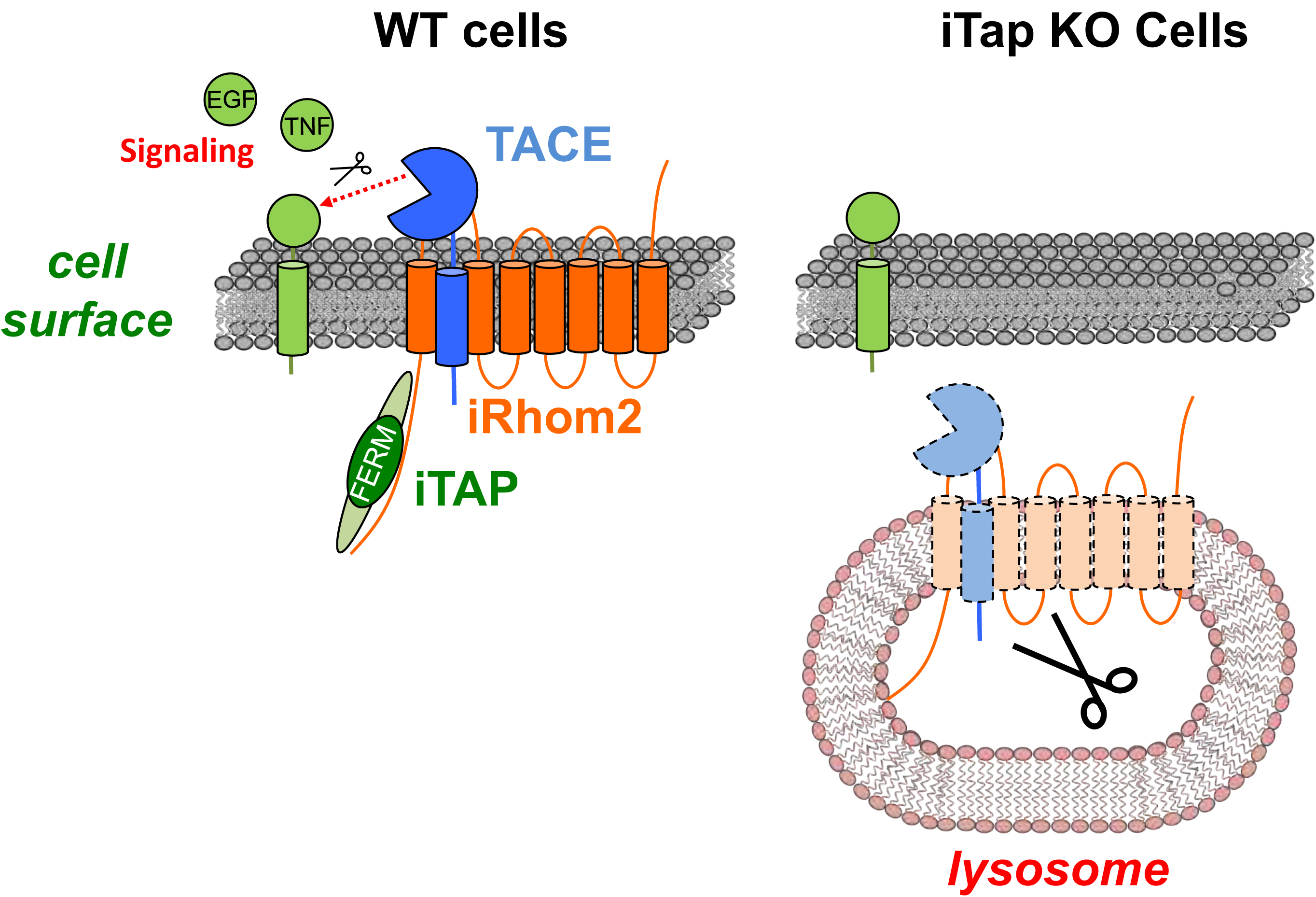
Model for iTAP function. Schematic representation of a speculative model for iRhom/TACE fate in the presence or absence of iTAP. In the presence of iTAP, the levels of iRhom2 and mature TACE are stabilized on the plasma membrane, enabling TACE to cleave signaling molecules such as TNF or EGFR ligands. By contrast, in iTAP KO cells, iRhom2/mature TACE are degraded because of mis-sorting to the lysosome, preventing signaling.

**Figure S5.**
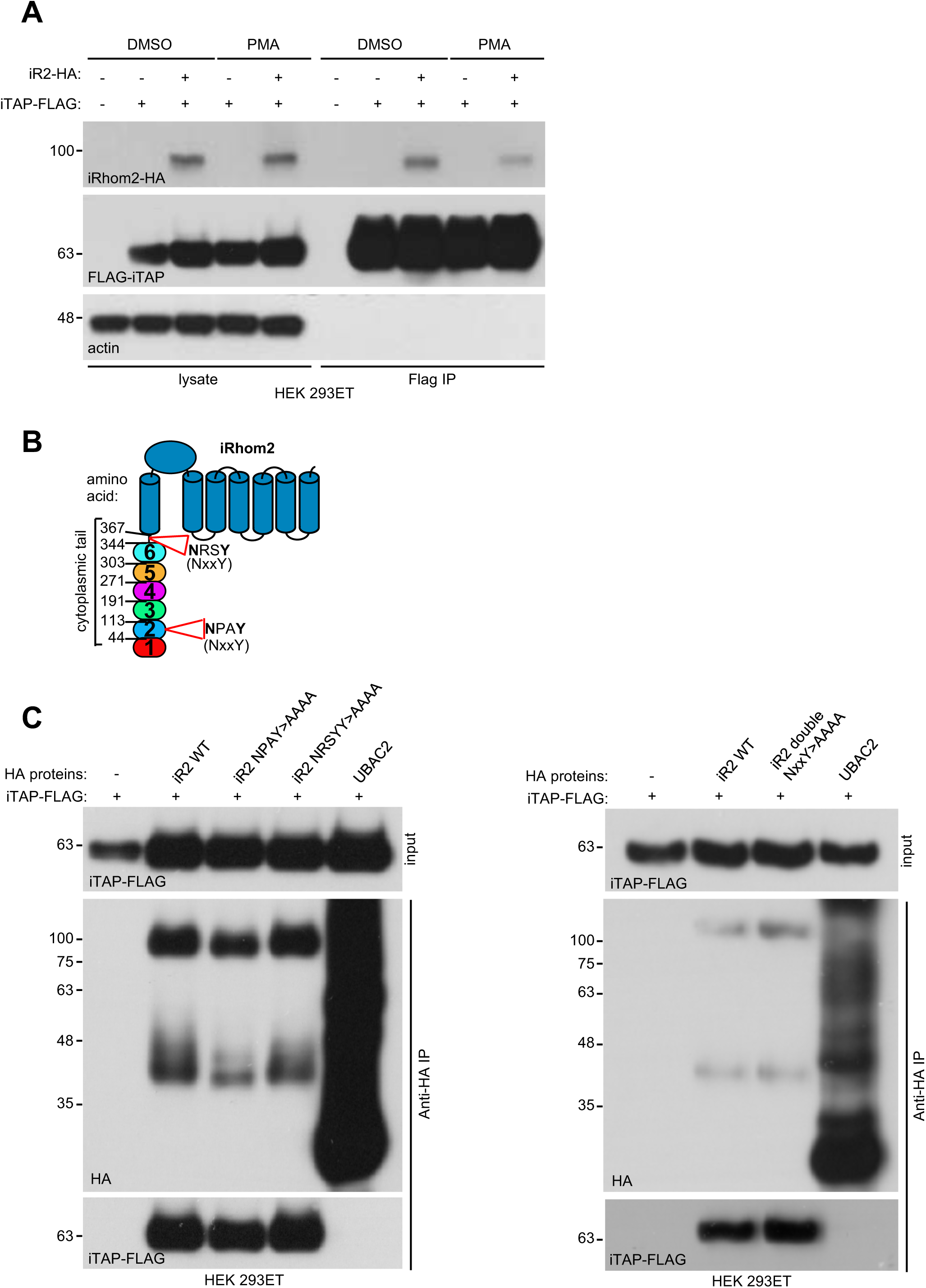
iTAP does not bind to features commonly recognized by FERM domain proteins. (**A**) iTAP appears not have a high affinity for actin. HEK 293ET cells transiently expressing vector or iRhom2-HA and iTAP-FLAG were starved overnight then stimulated with or without PMA (1μM) for 30min. PMA was shown to alter the steady state of iRhom interactions with clients (Cavadas et al., 2017) and hence is used to assess potentially differential binding upon stimulated conditions. An anti-FLAG IP was performed on the cell lysates and binding was assessed by western blotting (**B**) Schematic representation of the location of the NxxY motifs within the iRhom2 sequence. NxxY is a common consensus binding motif for FERM proteins (eg sorting nexins) that participate in vesicular sorting functions. (**C**). iTAP binding is independent of NxxY motifs. HEK 293ET cells were transiently transfected with iTAP-FLAG and vector, iRhom2-HA, single (left hand panel) or double (right hand panel) NxxY>AAAA mutants of iRhom2-HA, or Ubac2-HA as a negative control. PMA stimulation was performed as in S4A and cell lysates were subjected to anti-HA co-precipitation. iTAP binding to in iRhom immnoprecipitates was assessed by western blotting.

